# Extracellular vesicle cross-linking coupled with solid-phase purification facilitates discovery of ribonucleoprotein complexes assembled on EV RNAs

**DOI:** 10.64898/2026.10.03.756357

**Authors:** Rajat Dhyani, Stephen Clarke, Masashi Kuroda, Yohei Sanada, Marc Gorum, Bahnisikha Barman, Alissa Weaver, Takahisa Nakamura, Manuel Ascano

## Abstract

Extracellular vesicles (EVs) are fundamental mediators of intercellular communication that transfer macromolecular cargo, including proteins and RNAs, to functionally reprogram recipient cells. However, the molecular processes by which transferred ribonucleoprotein (RNP) complexes engage recipient host machinery to influence cellular behavior are poorly understood, primarily due to a dearth of methods capable of tracking these post-transfer interactions. To address this gap, we developed two complementary chemical biology platforms: EV-CLASP and EV(r)-CLASP. While EV-CLASP maps the native, intra-vesicular RNP landscape packaged within the EV, EV(r)-CLASP captures post-transfer dynamics by identifying the host proteins that directly associate with EV-derived RNAs following cellular uptake. Both approaches leverage metabolic labeling and photo-crosslinking for high-specificity recovery. Donor cells are cultured with 4-thiouridine (4SU) to biosynthetically label newly synthesized RNAs during EV biogenesis. Purified EVs can be crosslinked using 365 nm UV light to capture intra-vesicular RNP composition. Alternatively, 4SU-labeled EVs can be delivered to recipient cells and subsequently crosslinked to identify the primary physical interactions formed within the recipient-cell environment. Crosslinked RNP complexes are isolated under stringent conditions using solid-phase reversible beads to minimize background, followed by quantitative mass spectrometry for protein identification. Together, EV-CLASP and EV(r)-CLASP provide a robust, versatile framework for defining the dynamic RNP assemblies that facilitate extracellular RNA communication, offering a powerful toolkit to uncover novel RNA-associated regulatory networks, therapeutic targets, and biomarkers across diverse biological and disease contexts.

## Introduction

Extracellular vesicles (EVs) are heterogeneous, membrane-bound structures released by virtually all cell types to function as key mediators of local and systemic intercellular communication^1–3^. By transferring a diverse repertoire of bioactive cargo, including proteins, RNAs, lipids, metabolites, and DNA, EVs regulate numerous physiological and pathological processes, ranging from tissue development and immune modulation to cancer progression and metastatic dissemination^4,5^.

Accumulating evidence indicates that EV cargo loading is directed by active, regulated sorting mechanisms that selectively enrich discrete populations of RNAs, proteins, and other biomolecules, resulting in vesicle compositions that differ substantially from those of the parental cell^2^. RNA-binding proteins (RBPs) serve as critical arbiters of this selective trafficking, governing the packaging, stabilization, and secretion of specific transcript species into the vesicular lumen^6^. Upon internalization by recipient cells, these transferred RNAs can potentially reprogram recipient cells, altering their gene expression^6^. Despite substantial advances in EV cargo profiling, the molecular mechanisms that regulate RNA selection, packaging, and function within EV-mediated communication are poorly understood^2,3^. Indeed, how EV-delivered RNA functions in recipient cells remains largely a mystery, and it is unknown how that RNA engages the recipient cell machinery. Consequently, methods that enable the identification of EV-RNA-RBP interactions both within EVs and after they are delivered could reveal new therapeutic targets and mechanisms to engineer new drugs.

Conventional RNA interactome methodologies are powerful tools for studying RNA-protein interactions but typically require prior knowledge of specific RNA sequences or protein targets. Consequently, their utility for the systematic and unbiased characterization of the diverse repertoire of EV-associated RNA species, which includes mRNAs, microRNAs, long non-coding RNAs, and other regulatory transcripts, is limited. Furthermore, target-dependent capture strategies based on sequence-specific hybridization can be vulnerable to probe-mediated off-target interactions, potentially resulting in the enrichment of unintended nucleic acid species and reduced interaction specificity^7^. In addition, sequence-dependent hybridization approaches rely on open RNA regions for selective capture, which themselves represent a dynamic landscape, particularly for EV-derived RNAs in recipient cells. Because these metrics cannot readily discern between differential recovery arising from more efficient probe hybridization versus biological differences in RNA-protein interactions, such hybridization approaches introduce an inherent bias into the recovery of crosslinked ribonucleoprotein (RNP) complexes, confounding downstream protein enrichment analyses. Collectively, these limitations highlight the need for a sequence-independent strategy capable of capturing RNA-protein interactions across multiple stages of EV biology, including both RNA packaging within donor-derived EVs and the subsequent engagement of EV-transferred RNAs with recipient-cell proteins following uptake.

### Development of the protocol

To address this need, we developed Extracellular Vesicle-derived RNA–CrossLinking And Solid-phase Purification (EV-CLASP) as a sequence-independent platform that leverages metabolic incorporation of 4-thiouridine (4SU) that enables efficient UV-induced covalent crosslinking between RNAs and directly interacting proteins. Because the molecular cargo of EV is determined during EV biogenesis, perturbations that alter RNA trafficking, sorting, or packaging into EV can reshape both the RNA and protein composition of the vesicles. Consequently, EV-CLASP provides a powerful approach to systematically define the RNP associated with EV-derived RNAs within isolated large extracellular vesicles (LEV) or small extracellular vesicles (SEV), thereby providing mechanistic insight into how donor-cell biology regulates EV cargo composition.

However, EV biology extends beyond cargo packaging, as the functional consequences of EV-mediated communication are ultimately determined by the interactions that occur after EV uptake by recipient cells. Following internalization, EV-transferred RNAs engage a distinct set of recipient-cell proteins to form new RNP complexes that regulate downstream cellular responses. To establish a proof of concept these recipient-cell RNA-protein interactions can be selectively identified, we developed EV(r)-CLASP (where *r* denotes recipient), an extension of the EV-CLASP workflow that combines 4SU-labeled donor-cell RNAs with isotopically labeled recipient-cell proteins to selectively identify recipient proteins that directly associate with EVs-transferred RNAs following internalization.

In both platforms LEV (>200 nm) and SEV (<200 nm) containing labelled are isolated from conditional medium by differential or density gradient ultracentrifugation and subjected to multiple washing steps to remove residual extracellular 4SU, free RNA species, and soluble protein contaminants. Upon irradiation with 365 nm ultraviolet (UV) light, the thiol group of 4SU facilitates efficient covalent crosslinking between labeled RNAs and directly interacting proteins.

A key feature of the CLASP workflow is the use of Solid Phase Reversible Immobilization (SPRI) under highly denaturing conditions to selectively enrich covalently crosslinked RNA-protein complexes. Unlike affinity or sequence based purification methods, SPRI beads provide a sequence-independent means of capturing RNA. Following UV crosslinking, EVs are lysed in a buffer containing sodium dodecyl sulfate (SDS) and heated to disrupt membranes and dissociate noncovalent protein-protein and protein–RNA interactions. Under denaturing conditions, total nucleic acids are efficiently immobilized on SPRI magnetic beads, whereas only proteins covalently crosslinked to RNA remain associated during purification. Multiple stringent washing steps are performed under denaturing conditions to remove nonspecific contaminants, free proteins, membrane-associated impurities, and residual EV isolation artifacts. Crosslinked RNA-protein complexes are subsequently eluted from the beads, and RNA is digested using Benzonase nuclease to release crosslinked proteins for downstream analysis.

Label-free quantitative proteomics provides a straightforward and robust approach for identifying RNA-binding proteins enriched by EV-CLASP, as all proteins recovered originate from the isolated extracellular vesicles. In contrast, EV(r)-CLASP interrogates RNA-protein interactions that occur after 4SU labeled EV uptake by recipient cells, where proteins from both donor-derived EVs and recipient cells are present. Distinguishing the cellular origin of these proteins therefore requires a metabolic labeling strategy.To address this distinction, EV(r)-CLASP incorporates Stable Isotope Labeling by Amino Acids in Cell Culture (SILAC), in which recipient cells are cultured in heavy isotope-containing media while donor cells remain unlabeled. Following the uptake of 4SU-labeled EVs, recipient cells are irradiated with 365 nm UV light to induce covalent crosslinking between EV-transferred RNAs and interacting recipient-cell proteins. The resulting RNA-protein complexes are purified using the EV-CLASP workflow under highly denaturing conditions, ensuring the selective enrichment of covalently crosslinked complexes, and are subsequently identified and quantified by LC-MS/MS. By integrating orthogonal RNA labeling (4SU) with metabolic proteome labeling (SILAC), EV(r)-CLASP enables the selective identification of recipient-cell proteins that interact with EV-delivered RNPs.

### Applications of the method

EV-CLASP and EV(r)-CLASP provide versatile and scalable platforms for the comprehensive investigation of EV-associated RNP complexes and their functional roles in intercellular communication. By integrating metabolic RNA labeling, UV crosslinking, sequence-independent solid-phase purification, and mass spectrometry-based proteomics^8,9,10^, these methodologies enable unbiased and proteome-wide characterization of RBPs associated with EV cargo and recipient-cell remodeling processes.

A primary application of EV-CLASP is the global identification of EV-associated RNA-binding proteins and ribonucleoprotein assemblies under physiological and pathological conditions. The method enables systematic characterization of proteins directly associated with EV-derived RNAs, including messenger RNAs, long noncoding RNAs, microRNAs, circular RNAs, and other regulatory RNA species. Because EV-CLASP preserves both stable and transient RNA-protein interactions through covalent crosslinking, the protocol is particularly valuable for capturing dynamic or low-affinity interactions that are often lost using conventional RNA interactome methods^11–13^. In addition, the sequence-independent nature of EV-CLASP minimizes biases associated with antibody dependent or probe-based enrichment strategies^14–16^, allowing broad and unbiased interrogation of EV-associated RNP landscapes.

EV-CLASP is highly applicable to studies investigating EV biology in cancer, neurodegeneration, immune signaling, and metabolic disorders^6,17,18^. In cancer biology, EV-CLASP can be used to identify tumor-derived EV-associated RBPs involved in metastasis, immune evasion, angiogenesis, and therapeutic resistance^17,18^. The approach can also facilitate the discovery of EV-associated biomarkers for disease diagnosis, prognosis, and treatment monitoring^19,20^. In infectious disease research, EV-CLASP can be used to study RNA-protein interactions within extracellular vesicles released during viral or bacterial infections, helping to reveal how pathogens communicate with host cells and influence immune responses. The method can also be applied to investigate EV-mediated communication in cancer, stem cell systems, and immune cell populations, providing insights into how EV-associated RNAs and proteins regulate cellular function and intercellular signaling.

While EV-CLASP characterizes the native state of the vesicle, EV(r)-CLASP is designed to capture the dynamic interactions following internalization. This extension allows for a high-resolution view of how EV-derived RNPs are remodeled and integrated into recipient-cell regulatory pathways. Through integration of orthogonal metabolic labeling and SILAC-based quantitative proteomics^21,22^, EV(r)-CLASP distinguishes donor-derived EV cargo from the recipient-cell-associated RNA-protein interactions following EV uptake. This capability provides unique opportunities to study the fate of transferred EV-associated RNP complexes and their contribution to recipient-cell reprogramming. EV(r)-CLASP is particularly useful for investigating mechanisms of EV-mediated signaling in intercellular communication. The method can be applied to determine how EV-associated RNAs and RBPs alter transcriptional, translational, and post-transcriptional regulatory pathways within recipient cells. EV(r)-CLASP can be used to characterize how tumor-derived EVs reprogram stromal cells, immune cells, and metastatic niches through RNA-protein transfer. In addition, the platform can facilitate the investigation of neuron–glia communication and the EV-mediated propagation of neurodegeneration-associated RNA-binding proteins.

Importantly, EV(r)-CLASP enables temporal analysis of EV-mediated RNA regulatory remodeling by monitoring recipient-cell-associated RNA-protein interactions at different time points after EV internalization. Together, EV-CLASP and EV(r)-CLASP establish complementary methodologies for systematic and quantitative analysis of EV-associated RNA-protein interactions, EV cargo transfer, and recipient-cell functional reprogramming across diverse biological systems and disease contexts.

We optimized and implemented EV-CLASP and EV(r)-CLASP in colorectal cancer models using donor and recipient cell lines, including KRAS-mutant DLD-1 and DKs-8 cells, to systematically investigate EV-mediated RNA-protein transfer and recipient cell functional response. EVs isolated from DLD-1 donor cells were analyzed using EV-CLASP to identify tumor-associated RBPs and RNP complexes enriched within EV cargo. Following uptake of DLD-1-derived EVs by DKs-8 recipient cells, EV(r)-CLASP was used to identify recipient cell proteins that interact with LEV-delivered RNAs. Application of this protocol led to the identification of hundreds of shared and unique LEV-associated proteins, while also revealing extensive LEV-mediated alterations in recipient cell RNA-binding protein complexes. Collectively, these findings highlight the power of EV-CLASP and EV(r)-CLASP as robust platforms for dissecting dynamic EV-associated RNP interactions and elucidating their functional roles in colorectal cancer progression and intercellular communication.

### Comparison to other methods

EV-CLASP and EV(r)-CLASP differ from existing RNA interactome methodologies, including RNA Immunoprecipitation (RIP)^23,24^, Crosslinking and Immunoprecipitation (CLIP)^25^ and enhanced CLIP (eCLIP)^26^, Chromatin Isolation by RNA Purification-Mass Spectrometry (ChIRP-MS)^27^, and RNA Antisense Purification-Mass Spectrometry (RAP-MS)^28^, in two fundamental ways: first, EV-CLASP selectively identifies proteins associated with metabolically labeled EV-derived RNAs without requiring prior knowledge of RNA sequence or protein targets. Second, EV-CLASP employs a sequence-independent purification strategy under highly denaturing conditions, enabling enrichment of covalently crosslinked RNA-protein complexes while minimizing recovery of non-covalent or indirectly associated proteins. As a result, EV-CLASP can be used to characterize RNA-protein interactions associated with heterogeneous EV RNA populations, including RNA species that are incompletely characterized or not readily accessible to sequence-targeted approaches.

Global RNA interactome approaches such as OOPS and RIC provide broader, more sequence-independent profiling of RNA-binding proteins^11,12,29,30^; however, each method has inherent technical and biochemical limitations that may complicate the analysis of extracellular vesicle-associated ribonucleoprotein complexes. In OOPS, for example, proteins, lipids, and other biomolecules can nonspecifically partition into the interphase during organic phase separation together with crosslinked RNA-protein complexes, thereby complicating downstream proteomic analyses^29,30^. Likewise, RIC primarily enriches proteins associated with polyadenylated RNAs through oligo(dT)-based capture, potentially excluding non-polyadenylated EV-associated RNA species, including microRNAs, circular RNAs, and several regulatory noncoding RNAs^11,12^. In contrast, EV-CLASP and EV(r)-CLASP leverage metabolic 4SU labeling to selectively mark RNAs destined for EV incorporation. Subsequent UV-induced crosslinking at 365 nm generates covalent bonds between labeled RNAs and directly interacting proteins, enabling selective enrichment of EV-derived RNA-protein complexes under highly stringent purification conditions.

A distinguishing feature of EV-CLASP and EV(r)-CLASP is the use of paramagnetic bead-mediated solid-phase purification under highly denaturing conditions. Compared with chemically driven liquid–liquid phase separation or silica-based nucleic acid purification approaches^11,12,29,30^, magnetic bead-based enrichment enables more efficient, selective, and reproducible recovery of crosslinked RNA-protein complexes while substantially reducing nonspecific contaminants and sample loss^31–33^. Importantly, the protocol supports stringent washing under highly denaturing conditions, thereby preserving only covalently stabilized RNA-protein interactions and significantly improving purification specificity and downstream mass spectrometry sensitivity. Furthermore, EV(r)-CLASP extends these advantages through integration of orthogonal metabolic labeling and SILAC-based quantitative proteomics^21,34^, enabling discrimination between donor-derived EV cargo and recipient cell-associated RNA-protein remodeling following EV uptake. Collectively, EV-CLASP and EV(r)-CLASP establish robust, scalable, and highly sensitive platforms for comprehensive analysis of EV-associated RNA-protein interactions and EV-mediated intercellular communication.

### Experimental design

#### EV and host cell propagation

Donor and recipient cell lines used for EV-CLASP and EV(r)-CLASP can be selected according to the biological question, disease model, and EV system under investigation.

Because the efficiency of crosslinked RNA-protein complex recovery depends largely on the abundance of 4SU-labeled EV-associated RNAs secreted by donor cells, robust incorporation of 4SU into newly synthesized donor-cell RNAs and efficient EV production are critical for obtaining high-quality proteomic signals. Donor cells with high transcriptional activity, strong EV secretion capacity, and efficient metabolic uptake of 4SU generally provide improved labeling efficiency and increased recovery of EV-associated RNP complexes. Efficient intracellular uptake of 4SU requires expression of equilibrate nucleoside transporters, particularly ENT1 (SLC29A1)^35^ and related transporter family members. Therefore, optimizing 4SU concentration, labeling duration, and transporter expression levels in donor cells can substantially improve incorporation efficiency and downstream crosslinking sensitivity^36^. However, excessive 4SU exposure may affect cell viability, RNA metabolism, or EV biogenesis, and therefore labeling conditions should be empirically optimized for each cellular system.

#### Optimization and Validation of 4SU Labeling

To determine a 4SU concentration that supports efficient RNA labeling while maintaining cell viability, DLD-1 cells were seeded in 12-well plates at a density of 2.5 × 10^5^ cells per well and cultured for 24 h at 37 °C in a humidified incubator with 5% CO_2_. Following cell attachment, the culture medium was replaced with DMEM containing 4SU at final concentrations of 25, 50, 100, 150, or 200 μM. Vehicle control cells received an equivalent volume of DMSO. Cells were incubated for an additional 48 h, after which cell viability was assessed by trypan blue exclusion. Briefly, cells were washed with phosphate-buffered saline (PBS), detached by trypsinization, and resuspended in culture medium to generate a single-cell suspension. An aliquot of the cell suspension (10 μL) was mixed with an equal volume of 0.4% trypan blue solution and loaded onto a Countess II Automated Cell Counter chamber slide. Live and dead cells were quantified using a Countess II Automated Cell Counter. Viable cells were identified as trypan blue-negative cells, whereas non-viable cells were identified as trypan blue-positive cells. Percent cell viability was calculated and plotted (Supplementary Fig. 1a).

To determine the efficiency of 4SU incorporation into EV-associated RNAs, DLD-1 cells were cultured in the presence of 100 μM 4SU for 48 h under standard growth conditions. Conditioned medium was collected, and EVs were isolated by differential ultracentrifugation (Fig. 1). Purified EV were exposed to 365 nm UV light to induce covalent crosslinking between 4SU-labeled RNAs and associated proteins. Total RNA was subsequently isolated using TRIzol reagent according to the manufacturer’s instructions. RNA concentration and purity were assessed prior to downstream analysis. Purified RNA was enzymatically digested into individual ribonucleosides and analyzed by high-performance liquid chromatography (HPLC) as previously described^9^ (Supplementary Fig. 1b). The efficiency of 4SU incorporation was determined by quantifying the abundance of 4SU relative to total uridine content and calculated as:

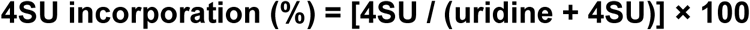

**Fig. 1.**
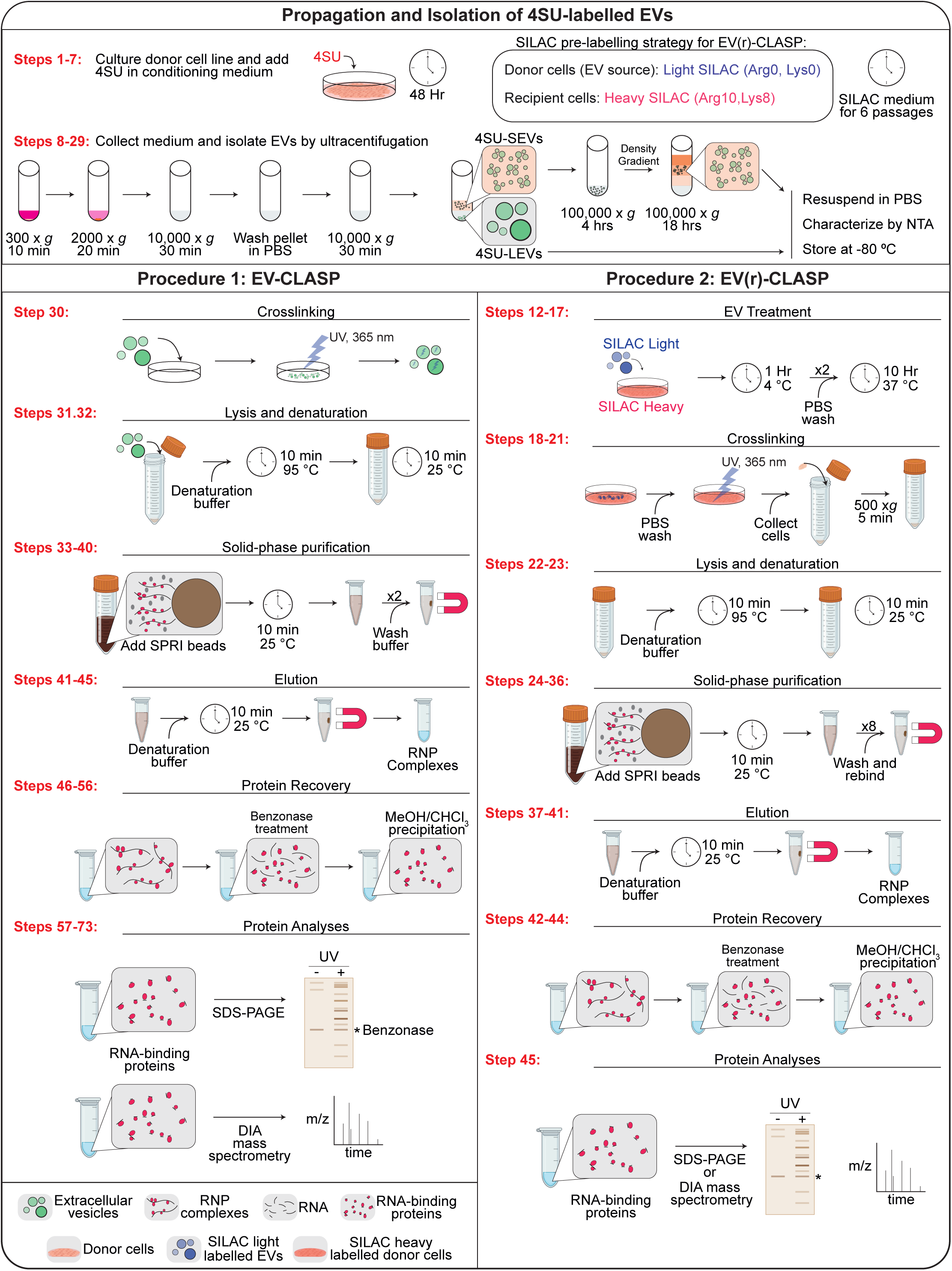
Outline of EV-CLASP and EV(r)-CLASP. The EV-CLASP and EV(r)-CLASP protocols begin with metabolic labelling of donor cells with 4-thiouridine (4SU) to label EV-associated RNAs. For EV(r)-CLASP, donor cells are additionally labelled with SILAC amino acids to enable quantitative discrimination of donor- and recipient-derived proteins (Steps 1-7). EVs are isolated from conditioned medium by differential ultracentrifugation (Steps 8-29) and characterized to assess their quality and purity. In EV-CLASP, purified EVs are subsequently irradiated with 365 nm UV light to induce covalent crosslinking between 4SU-labeled RNAs and associated proteins, thereby stabilizing EV-associated RNA–protein interactions (Step 30). Crosslinked EVs are lysed under highly denaturing conditions, and RNA–protein complexes are enriched using SPRI-based magnetic bead purification (Steps 31-40). Following stringent washing, RNA-bound proteins are eluted using 1× denaturation buffer (Steps 41-45). Residual RNA is then digested with Benzonase, and proteins are recovered by methanol/chloroform precipitation prior to LC– MS/MS analysis for the identification of EV-associated RNA-binding proteins and ribonucleoprotein complexes (Steps 46-73). In EV(r)-CLASP, SILAC-labeled EVs generated from donor cells cultured in light medium are incubated with recipient cells maintained in heavy SILAC medium to investigate EV-mediated RNA transfer and the remodeling of RNA–protein interaction networks in recipient cells (Steps 12–17). Following EV uptake, recipient cells are exposed to 365 nm UV light to induce covalent crosslinking between transferred 4SU-labeled EV RNAs and their interacting proteins, thereby capturing RNA–protein interactions formed after EV delivery (Steps 18–21). Cells are then lysed under highly denaturing conditions, and crosslinked RNA–protein complexes are enriched using SPRI-based magnetic bead purification (Steps 22–36). Following stringent washing, RNA-bound proteins are eluted with 1× denaturation buffer (Steps 37–41). Residual RNA is subsequently degraded by Benzonase treatment, and proteins are recovered by methanol/chloroform precipitation before being subjected to LC–MS/MS analysis for the identification and quantification of proteins associated with EV-derived RNAs in recipient cells (Steps 42-58).

#### Optimization of cross-linking conditions

Successful implementation of EV-CLASP and EV(r)-CLASP depends on efficient UV-induced crosslinking of 4SU-labeled RNAs to interacting proteins following EV purification or EV uptake by recipient cells, respectively. Irradiation with 365 nm UV light should be optimized to maximize crosslinking efficiency while minimizing RNA degradation or protein damage. Because only covalently stabilized RNA-protein complexes survive downstream denaturing purification conditions, efficient crosslinking is essential for preserving both stable and transient RNA-protein interactions and ensuring sensitive proteome-wide identification of EV-associated RNA-binding proteins.

### Optimization of CLASP bead buffer conditions

The SPRI method employs carboxyl-coated magnetic beads that reversibly bind RNA in SPRI bead buffer containing 20% (wt/vol) polyethylene glycol (PEG) and 2.5 M NaCl^10^. Increasing the ratio of bead buffer volume to sample volume (1X denaturation buffer) can enhance the recovery efficiency of smaller RNA species. In the EV-CLASP and EV(r)CLASP protocols, SPRI beads are used under protein-denaturing conditions to purify crosslinked RNA–protein complexes associated with EVs or recipient cells. Similarly, the ratio of bead buffer to sample buffer can influence the recovery efficiency of smaller RNAs. Although increasing this ratio may improve purification of smaller RNA molecules, excessively high bead buffer concentrations may not be optimal under CLASP conditions because denaturing agents present in the sample buffer can create PEG-associated solubility issues. Therefore, the CLASP bead buffer-to-sample buffer ratio should be optimized based on the experimental system and the size distribution of the RNAs of interest. For the EV-CLASP and EV(r)-CLASP protocols, SPRI bead buffer conditions were optimized to maximize the recovery of EV-associated RNAs together with crosslinked RNA-binding proteins for downstream quantitative proteomic analysis.

#### Verification of complete isotopic incorporation of SILAC amino acids

The extent of isotopic incorporation of SILAC-labelled amino acids was assessed to confirm complete labelling, as previously described^34^.

#### Skills and expertise necessary for protocol execution

Implementation of EV-CLASP and EV(r)-CLASP can be performed by a trained graduate student or postdoctoral researcher with experience in molecular biology, extracellular vesicle isolation, and proteomics. Investigators should have appropriate training in cell culture, handling biological samples, UV-crosslinking procedures, and biochemical purification methods. Access to proteomics core facilities and expertise in LC–MS/MS operation and data analysis are also required for successful execution of the protocol.

#### Limitations

Although EV-CLASP and EV(r)-CLASP provide scalable and systematic approaches for identifying EV-associated RNA-protein interactions, several experimental considerations should be carefully evaluated during implementation. A major determinant of the success of the protocol is the biological source of extracellular vesicles and the match to an appropriate recipient cell line. EV yield and composition vary substantially across cell types, physiological states, culture conditions, and EV subtypes differing in abundance, molecular content, and recovery efficiency. The protocol described here was optimized using colorectal cancer cell lines; however, adaptation to other cell types may require optimization of EV production, isolation strategies, and experimental scale to ensure sufficient recovery of EV material for robust RNA–protein interactome profiling.

Both EV-CLASP and EV(r)-CLASP rely on efficient incorporation of 4-thiouridine (4SU) into donor-cell RNAs, followed by UV-induced crosslinking of RNA-protein complexes. Consequently, optimization of 4SU labeling conditions may be required depending on the cell type, labeling duration, and experimental system to achieve efficient crosslinking while minimizing cellular toxicity and preserving normal cellular physiology. Recovery of crosslinked proteins further depends on the abundance of 4SU-labeled EV-associated RNAs, the efficiency of EV production and purification, and, in the case of EV(r)-CLASP, efficient uptake and internalization of EVs by recipient cells.

The quality of EV preparations also represents a critical factor influencing experimental outcomes. Isolation approaches based on differential centrifugation or ultracentrifugation may co-isolate protein aggregates, lipoproteins, membrane fragments, and other extracellular contaminants. Therefore, rigorous purification procedures and appropriate EV characterization should be implemented in accordance with the Minimal Information for Studies of Extracellular Vesicles (MISEV) guidelines established by the International Society for Extracellular Vesicles (ISEV) to ensure reproducibility and minimize background signals^37^. Because EV-associated RNA-protein complexes are often present at low abundance, careful optimization of UV-crosslinking efficiency, denaturing lysis conditions, and SPRI-based purification parameters is essential to maximize recovery and specificity.

As with other proteome-wide interaction mapping approaches, candidate proteins identified by EV-CLASP and EV(r)-CLASP should be independently validated using complementary biochemical, molecular, imaging, or functional assays. Validation under physiologically relevant EV exposure conditions is particularly important for confirming the biological significance of identified RNA-protein interactions and recipient-cell remodeling events. Finally, successful implementation of these protocols requires access or establishment of a quantitative proteomics infrastructure, high-resolution LC–MS/MS instrumentation, and appropriate computational resources for large-scale proteomic and statistical analyses. Several experimental parameters, including 4SU labeling duration, UV-crosslinking dosage, EV isolation conditions, SPRI bead-to-sample ratios, denaturing buffer composition, and SILAC labeling strategies, may require empirical optimization depending on the biological model system and experimental objectives.

### Materials

#### Biological materials

!CAUTION When working with colorectal cancer cell lines, several experimental precautions should be considered to ensure reproducibility, cell line integrity, and biological relevance. Cell lines should be routinely authenticated using short tandem repeat (STR) profiling and regularly tested for mycoplasma contamination, as prolonged culturing and cross-contamination can significantly alter cellular behavior and experimental outcomes^38,39^. Culture conditions, including serum source, cell density, and media composition, should be standardized across experiments. For extracellular vesicle studies, cells should preferably be maintained in media supplemented with dialyzed fetal bovine serum (FBS) or EV-depleted serum to minimize contamination from serum-derived extracellular vesicles^40^, free amino acids, and small metabolites that may interfere with EV isolation, SILAC labeling efficiency, and downstream proteomic analyses.

!CAUTION EV samples and cultured cells should be handled in accordance with approved institutional biosafety guidelines and laboratory safety regulations. Proper handling and disposal of biological samples are essential to minimize the risk of exposure to potentially bioactive or pathogenic materials and to prevent sample contamination during EV-CLASP and EV(r)-CLASP protocols.

##### Reagents

- DMEM (Gibco, cat. no. 11965-118)
- L-Glutamine (Gibco, cat. no. A2916801)
- Sodium pyruvate (Gibco, cat. no. 11360070)
- MEM non-essential amino acids solution (Gibco, cat. no. 11140050)
- 4-Thiouridine (Sigma-Aldrich, cat. no. T4509)
- DMSO (Sigma-Aldrich, cat. no. D2650)
- PBS (National Diagnostics, cat. no. CL-253)
- FBS (Peak Serum, cat. no. PS-FB2)
- Tris-base (Sigma-Aldrich, cat. no. 93362)
- HCl (Sigma-Aldrich, cat. no. H1758)
- NaCl (Sigma-Aldrich, cat. no. S3014)
- EDTA (Sigma-Aldrich, cat. no. E6758)
- NaOH (Sigma-Aldrich, cat. no. S8045)
- Sucrose (Sigma-Aldrich, cat. no. 84097)
- SDS (Sigma-Aldrich, cat. no. 74255)
- IGEPAL^®^ CA (Thermo Scientific, cat. no. J61055.AP)
- Glycerol (Sigma-Aldrich, cat. no. G5516)
- Sera-mag Speed Beads (Thermo Fisher Scientific, cat. no. 09-981-123)
- Tween 20 (Sigma-Aldrich, cat. no. P9416)
- PEG-8000 (Sigma-Aldrich, cat. no. 89510)
- Benzonase (Novagen, cat. no. 70746)
- MgCl_2_ (Sigma-Aldrich, cat. no. M8266)
- DTT (Sigma-Aldrich, cat. no. D9779)
- Methanol (Sigma-Aldrich, cat. no. 34860)
- Chloroform (Sigma-Aldrich, cat. no. C2432)
- NuPAGE LDS Sample Buffer (4X) (Invitrogen, cat. no. NP0007)
- NuPAGE Sample Reducing Agent (10X) (Invitrogen, cat. no. NP0004)
- NuPAGE 10%, Bis-Tris, 1.0 mm, Mini Protein Gel, 10-well (Invitrogen, cat. no. NP0301BOX)
- NuPAGE MOPS SDS Running Buffer (20X) (Invitrogen, cat. no. NP0001)
- SimplyBlue Safe Stain (Invitrogen, cat. no. LC6065)
- Pierce SILAC Protein Quantification Kit (Trypsin)-DMEM (Thermo-Scientific A33972)
- OptiPrep Density Gradient Medium (Millipore Sigma, cat no. D1556)
- Pipetting Needles Blunt End Standard (Cadence Science, cat no. 7942)

##### Equipment

- Floor Low Speed Centrifuge (Thermo Fisher Scientific, cat. no. Sorvall RC-3B Plus)
- H-6000A 6 × 1000 mL Swinging-Bucket Rotor (Thermo Fisher Scientific, cat. no. 11250)
- H-6000A bucket adapter for 500 mL tubes (Thermo Fisher Scientific, cat. no. 00-444)
- Ultracentrifuge (Beckman coulter, cat. no. Optima LE-80k)
- Ti45 ultracentrifuge tubes (Beckman Coulter Cat #355622)
- SW 32 Ti Swinging-Bucket Rotor (Beckman coulter, cat. no. 369650)
- Spectrolinker XL-1500 (Spectronics, cat. no. XL-1500A)
- DynaMag-2 Magnet (Thermo Fisher Scientific, cat. no. 12321D)
- Microcentrifuges (Eppendorf, cat. no. 5404000138)
- Vortex-Genie 2 (Scientific Industries, cat. no. SI-0236)
- Digital Dry Bath (Thermo Fisher Scientific, cat. no. 11-715-125D)
- 3 mLultracentrifuge tubes–Beckman Coulter Cat #362305
- Ti45 ultracentrifuge tubes–Beckman Coulter Cat #355622
- SW40 tubes–Beckman Coulter Cat #344060
- Concentrators–Millipore/Sigma Cat #UFC501024
- Long metal syringe – Cadence Sci Ref 7942

##### Plasticwares

- 150 mm TC-treated cell culture dish (Corning, cat. no. 353025)
- 500 mL PP Centrifuge tubes (Corning, cat. no. 431123)
- Cell scraper (VWR, cat. no. 101093-452)
- 1.5 mL tubes (Denville, cat. no. C2170)
- Thin wall polypropylene tube, 25 x 89mm (Beckman coulter, cat. no. 326823)
- 0.22 µm filter (Millipore, cat. no. S2GPU05RE)
- 0.45 µm filter (Millipore, cat. no. SCHVU05RE)
- Thick wall Polycarbonate tubes (Beckman Coulter 362305)
- Centrifuge Tubes (Beckman Coulter 344060)
- Centrifuge tubes (Beckman Coulter 344058)

##### Complete DMEM

Complete DMEM consists of DMEM supplemented with 2 mM L-glutamine, 1 mM sodium pyruvate, 1X MEM non-essential amino acids, and 10% (vol/vol) FBS. Store the complete medium at 4 °C for up to 3 months.

##### 4SU stock solution

Prepare a 1 M stock solution of 4SU in DMSO and aliquot into sterile 1.5 mL Eppendorf tubes. Store aliquots at -20 °C protected from light for up to 1 year.

**CRITICAL:** 4SU is highly light sensitive and should be handled under minimal light exposure.

##### 3X Denaturation Buffer

150 mM Tris-HCl (pH 6.8), 30% glycerol (v/v), 7.5% SDS (w/v), 2% IGEPAL^®^ CA

##### Bead buffer

2.5 M NaCl, 20% PEG-8000 (Dissolve overnight and make up to 500 mL with ddH_2_O).

##### 1X denaturation buffer

Add 10 mL 3X denaturation buffer to 20 mL ddH_2_O.

##### Wash buffer

To 20 mL ddH_2_O add 10 mL 3X denaturation buffer + 20 mL bead buffer.

##### Prepared SPRI Beads

Vortex until no clumps visible, add 1 mL Sera-mag Speed Beads (50 mg/mL) to tube and place on magnet. Carefully remove supernatant with pipette and wash with 1 mL TE buffer 3 times. Re-suspend in 1 mL TE buffer.

##### SPRI Bead suspension

18% (w/v) PEG-8000, 1 M NaCl, 10 mM Tris (pH 8.0), 1 mM EDTA, 0.055% Tween-20, 10 mM TE buffer, 1 mg/mL SPRI Beads. 4X Benzonase reaction buffer 80 mM Tris-HCl (pH 7.5), 20 mM MgCl_2_, 0.6 M NaCl, 4 mM DTT, 40% glycerol.

**CRITICAL** DTT should be added fresh before use.

##### SDS-PAGE sample buffer

NuPAGE 2X LDS with 1X sample reducing buffer.

##### SILAC Media preparation

Prepare trypsin-compatible SILAC media by dissolving 50 mg of ^13^C_6_ ^15^ N_2_ L-Lysine-2HCl (heavy) and 50mg of ^13^C_6_ ^15^N_4_ L-Arginine HCl (heavy) in 1 mL of media and mix thoroughly until fully dissolved. Add the dissolved amino acid mixture to a 500 mL bottle of media supplemented with dialyzed FBS and mix thoroughly. Similarly, to make a SILAC light media add 50 mg of L-Arginine-HCL and L-Lysine-HCL to a 500 mL bottle of media supplemented with dialyzed FBS and mix thoroughly.

#### Procedure 1: EV-CLASP for the systematic identification of extracellular vesicle-associated RNA-protein complexes

##### Propagation of 4SU-labeled EVs ● Timing ∼3 d

1. Seed 15 × 10^6^ DLD cells in 150 mm tissue culture dishes containing 20 mL complete DMEM media. Typically, 30 dishes are prepared per experimental condition to obtain sufficient material for downstream analyses. (Fig.1).

2. Incubate the cells overnight at 37 °C in a humidified incubator.

3. Verify that the cells have reached ∼80% confluency before proceeding.

**CRITICAL STEP** Carefully monitor cell confluency before conditioning. Over-confluent cultures may exhibit altered EV production and cellular stress responses, whereas under-confluent cultures may produce insufficient EV yields.

4. Remove the culture medium and wash the cells three times with PBS (10 mL per wash for 150 mm culture plates).

5. Add conditioning medium, preferably serum-free, using the same volume as the original culture medium.

6. Supplement the culture medium with 100 µM 4SU to generate labeled donor cell line EVs .

7. Incubate the cells for ∼48 h. The conditioning period may vary depending on the cell type, growth rate, and experimental objectives.

##### Isolation of 4SU labeled EVs ● Timing ∼ 8 h

8. Collect the conditioned medium from the flasks into 50 mL conical tubes, pooling samples for each condition. Before beginning EVs isolation, ensure that the ultracentrifuge is powered on, vacuum-locked, and pre-cooled to 4 °C.

9. Centrifuge the conditioned medium at 300 × g for 5 min to remove intact cells.

10. While samples are undergoing centrifugation, determine cell counts from at least one culture dish for every four dishes processed.

11. Transfer the supernatant to fresh 50 mL conical tubes and centrifuge at 2,000 × g for 25 min at 4 °C to remove residual cell debris.

12. Carefully transfer the resulting supernatant into Ti45 ultracentrifuge tubes and centrifuge at 10,000 × g for 30 min at 4 °C.

13. Ti45 ultracentrifuge tubes have a total capacity of 60 mL; therefore, ensure that each tube is filled to at least three-quarters of its volume.

14. The resulting pellet contains LEV. Carefully remove the supernatant containing SEV to a new 50 mL concial tube, and resuspend the pellet in 3 mL PBS, and transfer the suspension into 3 mL ultracentrifuge tubes.

**CRITICAL STEP:** When removing supernatants after centrifugation, avoid disturbing LEV pellets, which may be translucent and difficult to visualize.

15. Centrifuge the LEV samples at 10,000 × g for 30–60 min at 4 °C using a tabletop ultracentrifuge.

16. Carefully remove the supernatant and gently resuspend the LEV pellet in 100 μL PBS. Keep the LEV suspension on ice and perform nanoparticle tracking analysis (NTA) as soon as possible.

**CRITICAL STEP:** Perform all EV isolation steps at 4 °C whenever possible to minimize protein degradation and preserve EV integrity.

**PAUSE POINT**. Purified LEV pellets may be resuspended in PBS and stored at −80 °C for up to 1 month before downstream processing without detectable loss of sample quality. This storage step facilitates the collection of sufficient EV material from multiple preparations when required. Avoid repeated freeze–thaw cycles, as they may compromise EV integrity.

17. Wash the concentrators with 40-70 mL ddH2O and spin at 2,000×g for 2 minutes. Discard wash.

18. Transfer the supernatant containing SEV (obtained in Step 14) to centrifugal concentrators and centrifuge at 3,000 × g for 5 min at 4 °C using a tabletop centrifuge.

Repeat centrifugation as needed until the retentate volume is reduced to approximately 30 mL. Recover the concentrated supernatant from the concentrator and discard the flow-through.

19. Remove the concentrator from the collection tube, invert it onto a clean collection tube, and centrifuge at 3,000 × g for 5 min at 4 °C to recover the residual sample. Pool the recovered sample with the concentrated supernatant obtained in Step 19.

20. Cushion:

(A) Transfer the 30 mL concentrated supernatant to an SW32 ultracentrifuge tube.

(B) Carefully underlay the sample with 2 mL of 60% OptiPrep (iodixanol) using a long metal cannula or syringe, creating an iodixanol cushion at the bottom of the tube.

(C) Balance the SW32 tubes and ensure that each contains at least 32 mL of solution (30 mL supernatant plus 2 mL OptiPrep) to prevent tube collapse during ultracentrifugation. Centrifuge in an SW32 Ti swinging-bucket rotor at 100,000 × g for 4 h at 4 °C.

21. Using a long metal syringe, carefully collect the bottom 3 mL from each SW32 tube without disturbing the upper layers. Keep the fractions from each tube separate; do not pool them.

(A) Each 3 mL fraction consists of 2 mL of 60% OptiPrep (iodixanol) and approximately 1 mL of SEV that accumulate immediately above the iodixanol cushion.

(B) Mix each fraction thoroughly to obtain a homogeneous solution with a final OptiPrep concentration of 40% (vol/vol).

22. Prepare the OptiPrep (iodixanol) solutions required for the density gradient as shown in table 1

**Table 1.**
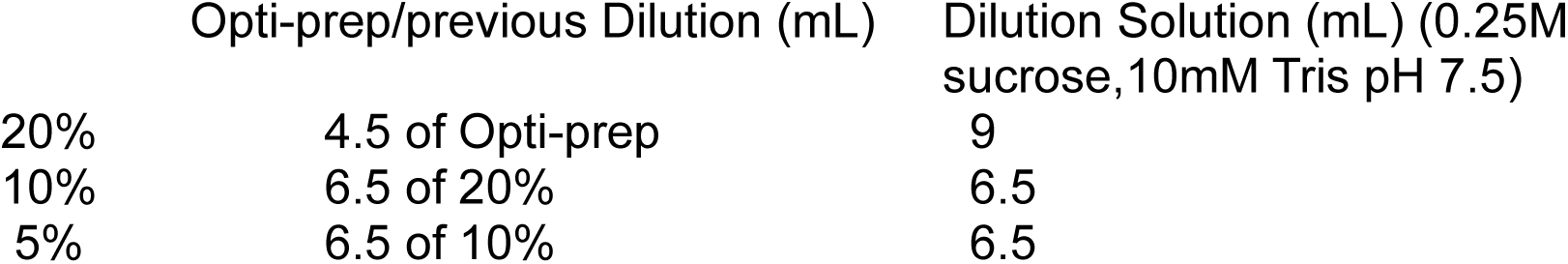
OptiPrep (iodixanol) solutions for density gradient preparation.

|  | Opti-prep/previous Dilution (mL) | Dilution Solution (mL) (0.25M sucrose, 10mM Tris pH 7.5) |
| --- | --- | --- |
| 20% | 4.5 of Opti-prep | 9 |
| 10% | 6.5 of 20% | 6.5 |
| 5% | 6.5 of 10% | 6.5 |

**Table 2.**
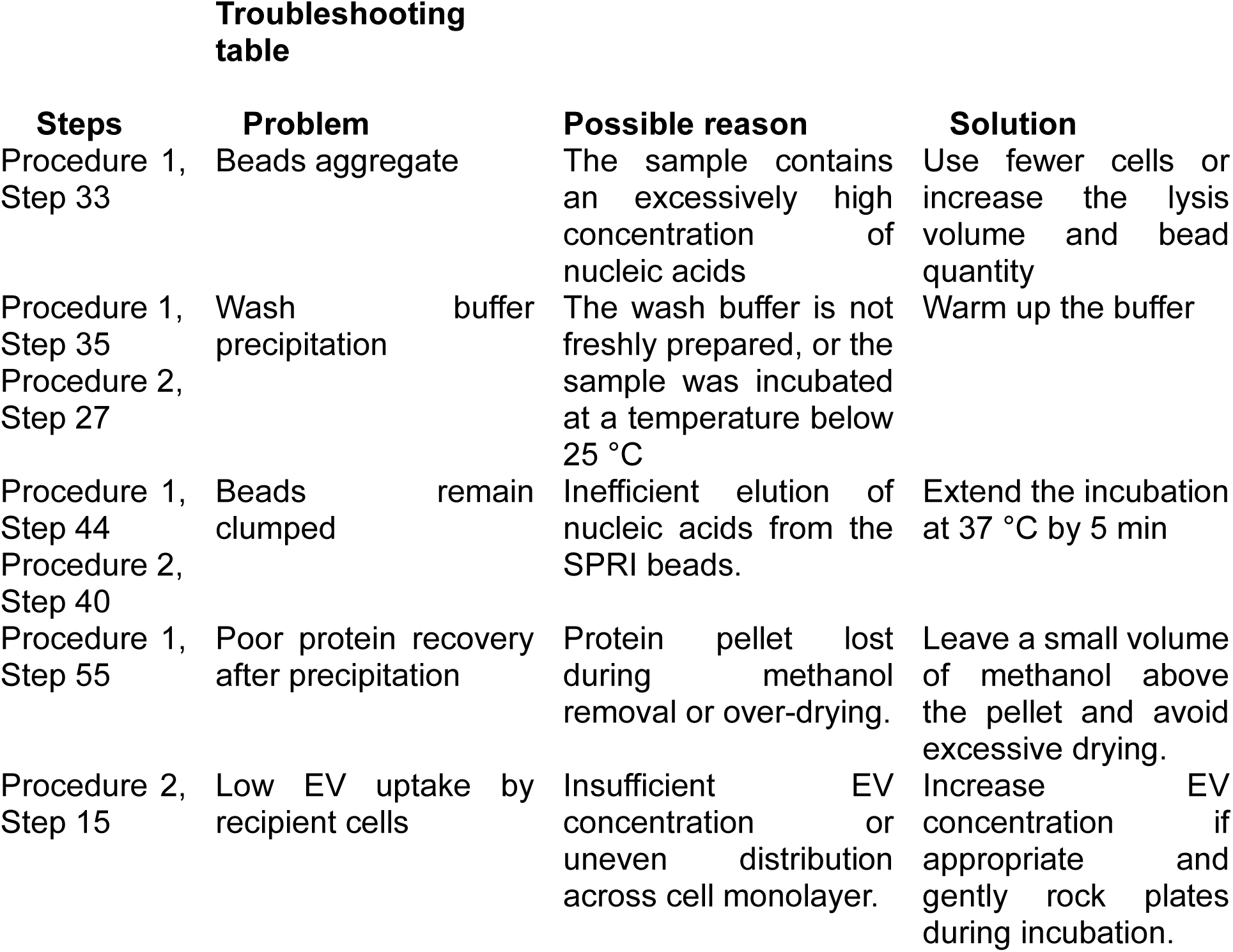
Troubleshooting table.

Troubleshooting Troubleshooting advice can be found on Table 2.
| Steps | Problem | Possible reason | Solution |
| --- | --- | --- | --- |
| Procedure 1, Step 33 | Beads aggregate | The sample contains an excessively high concentration of nucleic acids | Use fewer cells or increase the lysis volume and bead quantity |
| Procedure 1, Step 35<br>Procedure 2, Step 27 | Wash buffer precipitation | The wash buffer is not freshly prepared, or the sample was incubated at a temperature below 25 °C | Warm up the buffer |
| Procedure 1, Step 44<br>Procedure 2, Step 40 | Beads remain clumped | Inefficient elution of nucleic acids from the SPRI beads. | Extend the incubation at 37 °C by 5 min |
| Procedure 1, Step 55 | Poor protein recovery after precipitation | Protein pellet lost during methanol removal or over-drying. | Leave a small volume of methanol above the pellet and avoid excessive drying. |
| Procedure 2, Step 15 | Low EV uptake by recipient cells | Insufficient EV concentration or uneven distribution across cell monolayer. | Increase EV concentration if appropriate and gently rock plates during incubation. |

23. Prepare the 40/20/10/5% iodixanol density gradients. Transfer the 3 mL sample (adjusted to 40% iodixanol) to the bottom of an SW40 ultracentrifuge tube. Carefully overlay the sample with 3 mL of 20% OptiPrep (iodixanol), followed sequentially by 3 mL of 10% OptiPrep and 3 mL of 5% OptiPrep

**CRITICAL STEP:** Thoroughly remix each OptiPrep (iodixanol) solution immediately before use to ensure homogeneity. Carefully overlay each dilution onto the gradient slowly and gently to maintain discrete interfaces and prevent mixing between adjacent layers.

24. Centrifuge the density gradients in an SW40 Ti swinging-bucket rotor at 100,000 × g for 18 h at 4 °C using a floor ultracentrifuge.

25. Collect the 6th and 7th sequential 1 mL fractions of the density gradient into separate 3 mL microcentrifuge tubes, as these fractions are enriched for SEV.

26. Transfer each 1 mL fraction to a compatible ultracentrifuge tube and add 2 mL of PBS. Mix thoroughly, then ultracentrifuge in a TLA-110 rotor at 100,000 × g for 3 h at 4 °C using a tabletop ultracentrifuge.

27. **CRITICAL STEP:** The SEV pellet is often not visible following ultracentrifugation. Mark the outward-facing side of each tube before centrifugation and indicate the expected pellet location with a small circle to aid pellet recovery during supernatant removal.

28. Keep the fractions on ice and characterize the SEV using NTA as soon as possible.

**PAUSE POINT**: Purified SEV pellet can be stored at −80 °C for up to 1 month.

##### Characterization of EV by NTA and UV crosslinking ● Timing ∼2 h

29. Characterize LEV and SEV preparations for EV-CLASP and EV(r)-CLASP by NTA to determine particle concentration and size distribution (Box 1; Supplementary Figs. 2 and 3).

30. Prior to UV irradiation, divide each sample into two equal aliquots and transfer them into separate tubes. One aliquot should remain non-irradiated and serve as the negative control, while the other aliquot will be subjected to UV crosslinking. Expose the uncovered test sample dishes to 365 nm UV light twice, delivering a total energy dose of 1.2 J/cm², using a Spectrolinker XL-1500 or a comparable UV crosslinking system (Fig. 1). Retain the corresponding non-irradiated sample under identical conditions, except for UV exposure, to serve as a negative control.

**PAUSE POINT** Snap-freeze the EV preparations in liquid nitrogen and store at –80 °C until further processing. When ready to proceed with the CLASP protocol, resuspend the EVs directly in 1× denaturation buffer (∼900 µL) and mix thoroughly by pipetting up and down to ensure complete lysis.

#### EV-CLASP

##### Preparation of extracellular vesicle lysate and protein denaturation ● Timing ∼30 min

31. Incubate the samples in 900 µL 1X denaturation buffer from Step 30 at 95 °C for 10 min. Following boiling, transfer two 200 µL aliquots into 1.5 mL Eppendorf tubes as “input” samples. These input samples should be processed alongside EV-CLASP samples later at step 45 for RNA analysis and at step 59 for protein analysis.

**PAUSE POINT** EV lysates may be stored at -20 °C at this stage. To resume the protocol, thaw the samples and incubate at 95 °C for 10 min before proceeding to Step 32.

32. Incubate the samples at room temperature (RT) for 10 min.

##### Solid-phase purification ● Timing ∼3 h

33. Add 600 µL of SPRI bead suspension (see ‘Reagent setup’) to 900 µL of denatured lysate to result in a 2:3 bead buffer to sample ratio and mix thoroughly.

**?TROUBLESHOOTING**

34. Mix the samples by pipetting up and down 10 times, then incubate at 25 °C for 10 min. Place the tubes on a magnetic stand and allow the SPRI beads to collect on the magnet. Bead settling times may vary and can take up to 3 min.

35. Carefully remove and discard the supernatant, then add 1 mL wash buffer.

**?TROUBLESHOOTING**

36. Vortex the samples at maximum speed and allow the SPRI beads to re-pellet on the magnetic stand.

37. Remove and discard the supernatant. Resuspend the beads from one tube in 1 mL of wash buffer by vortexing at maximum speed. Transfer the bead suspension to the next tube and repeat the process sequentially to combine the contents of all four sample tubes into a single tube using the same 1 mL wash buffer suspension.

38. Place the tube containing the aggregated beads on a magnetic stand and allow the beads to fully collect.

39. Remove and discard the supernatant and replace it with 1 mL of wash buffer.

40. Vortex the beads at maximum speed to thoroughly resuspend them, then return the tube to the magnetic stand and allow the beads to fully re-pellet.

41. Remove the wash buffer supernatant and add 200 μL of 1× denaturation buffer. Thoroughly resuspend the beads by vortexing.

42. Briefly centrifuge the tubes at low speed for a few seconds to collect beads from the walls and caps of the tubes. Avoid over-centrifugation, as this may cause bead aggregation or clumping.

43. Incubate the samples at room temperature for 5 min.

44. Place the tubes on a magnetic stand to collect the beads and transfer the supernatant to a fresh tube. Save a 20 µL aliquot of the supernatant in a 1.5 mL eppendorf tube for Step 45.

**?TROUBLESHOOTING**

**PAUSE POINT** Eluted samples containing crosslinked RNA-protein complexes may be stored at -80 °C for several weeks. Avoid repeated freeze-thaw cycles, which may reduce sample quality and protein recovery.

45. (Optional) To assess RNA recovery by CLASP, treat the aliquot collected in step 60 with 1 mg/mL proteinase K for 2 h at 55 °C, followed by RNA purification using TRIzol according to the manufacturer’s instructions.

##### Benzonase treatment ● Timing ∼ 2 h

46. Add 250 µL of 4× Benzonase reaction buffer and 496 µL nuclease-free water containing 2.5 µL Benzonase to the 200 µL supernatant obtained from step 44.

47. Incubate the samples at 37 °C for 2 h.

##### Methanol and Chloroform Precipitation ● Timing: ∼45 min

48. Divide the 800 µL Benzonase-treated sample equally into two tubes (400 µL per tube).

49. Add an equal volume of 100% methanol (MeOH) to each sample to obtain a final concentration of 50% (vol/vol) MeOH. Vortex at maximum speed for 3s.

50. Add 200 µL chloroform to each tube and vortex at maximum speed for 3s.

51. Centrifuge the samples at 15,000 × g for 2 min at RT.

52. Carefully remove the upper aqueous/MeOH phase without disturbing the interphase.

**CRITICAL STEP** Note that the precipitated protein layer may not appear visibly white; therefore, caution should be taken to avoid disturbing the interface.

53. Add 400 µL MeOH to each tube and vortex at maximum speed for 10 s.

54. Centrifuge at 15,000 x g for 3 min at RT. The protein precipitate should form a pellet at the bottom of the tube.

55. Carefully remove as much MeOH as possible without disturbing the pellet, then briefly dry the pellets in a vacuum centrifuge for 5 min at RT.

**?TROUBLESHOOTING**

56. Resuspend each pellet in 20–50 µL of 0.5% SDS prepared in 100 mM Triethylammonium bicarbonate (TEAB) buffer. Pipette thoroughly up and down until the pellet is completely dissolved and a homogeneous solution is obtained.

**PAUSE POINT** Following Benzonase treatment and protein recovery, samples may be stored at −80 °C before protein digestion and LC–MS/MS analysis.

57. Add 10 µL of sample to 2X NuPAGE LDS Sample Buffer supplemented with 2X NuPAGE Sample Reducing Agent and mix thoroughly before heating and SDS–PAGE analysis.

**CRITICAL STEP**: The volume of 2× NuPAGE LDS Sample Buffer required for pellet re-suspension may vary depending on cell type, 4SU incorporation efficiency, crosslinking efficiency, and the extent of protein precipitation. Based on our experience, 50–70 µL is generally sufficient.

58. Incubate the samples at 95 °C for 5 min.

59. Visualize the precipitated LEV and SEV proteins by silver staining (Fig. 2a and Supplementary Fig. 2c) or continue directly to LC–MS/MS analysis^41^.

**Fig. 2.**
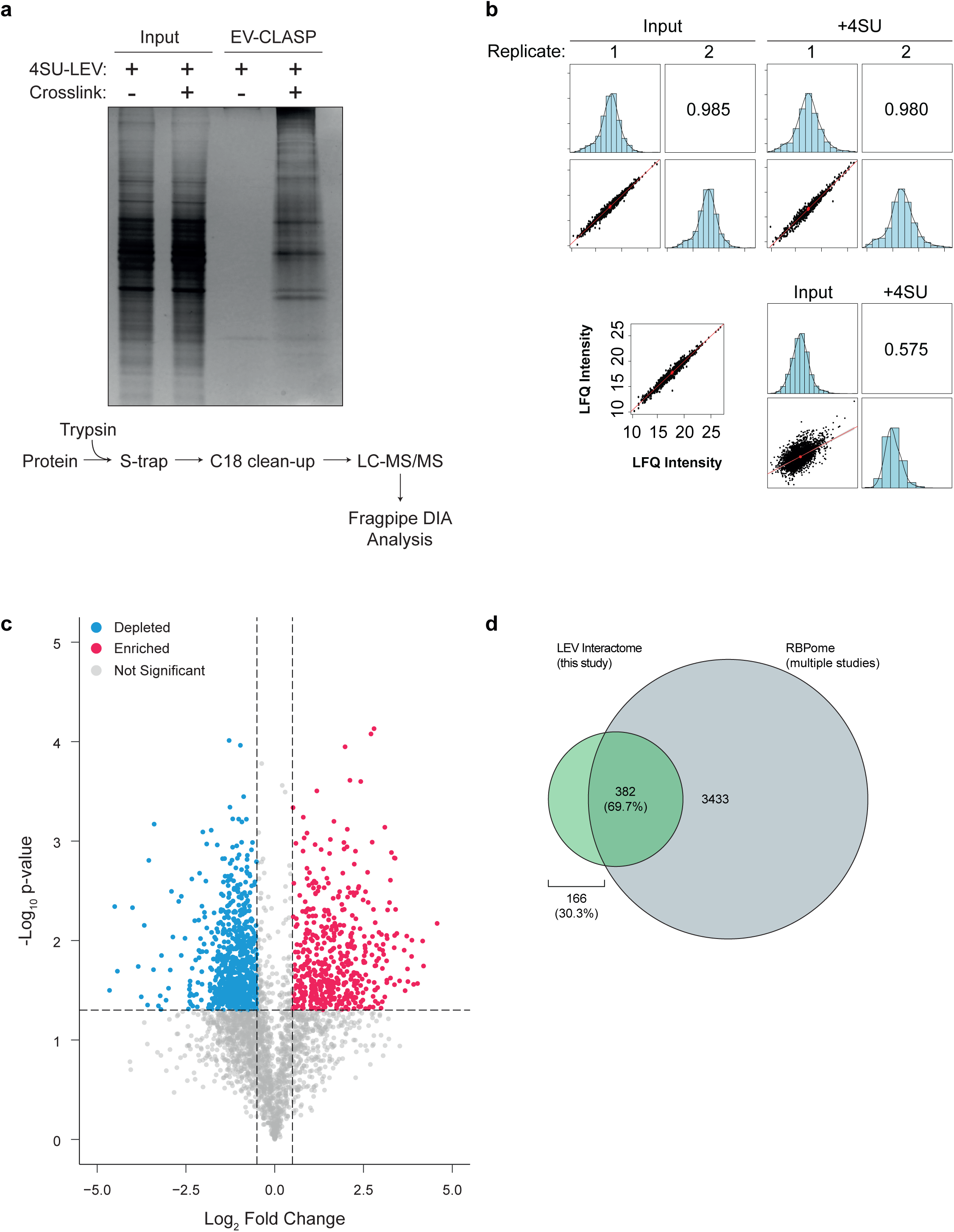
Anticipated results and proteomic analysis of EV-CLASP samples. **(a)** SDS– PAGE followed by silver staining of proteins enriched by EV-CLASP from 4SU-labeled extracellular vesicles (EVs) isolated from DLD colorectal cancer cell lines. **(b)** Pearson correlation coefficient analysis between all biological replicate samples of input and +4SU EV-CLASP conditions demonstrates high experimental reproducibility (top). Pearson correlation coefficient analysis demonstrates moderate correlation (r = 0.575) between LEV-Input and LEV-CLASP samples, highlighting distinct proteomic enrichment profiles associated with EV-CLASP purification (bottom). **(c)** Volcano plot showing differential protein enrichment in EV-CLASP samples relative to corresponding input controls. The x-axis represents log2 fold change, and the y-axis represents −log10 *P* value. Significantly enriched proteins identified in EV-CLASP samples are highlighted in red, whereas significantly depleted proteins are shown in blue. **(d)** The recovered protein set was compared with established RNA-binding protein (RBP) reference databases, such as RBP2GO, to evaluate enrichment of RNA-associated proteins. A substantial overlap with annotated RBPs indicates successful recovery of RNA-binding proteins and supports the capture of RNA-associated regulatory networks by the EV-CLASP

**Fig. 3.**
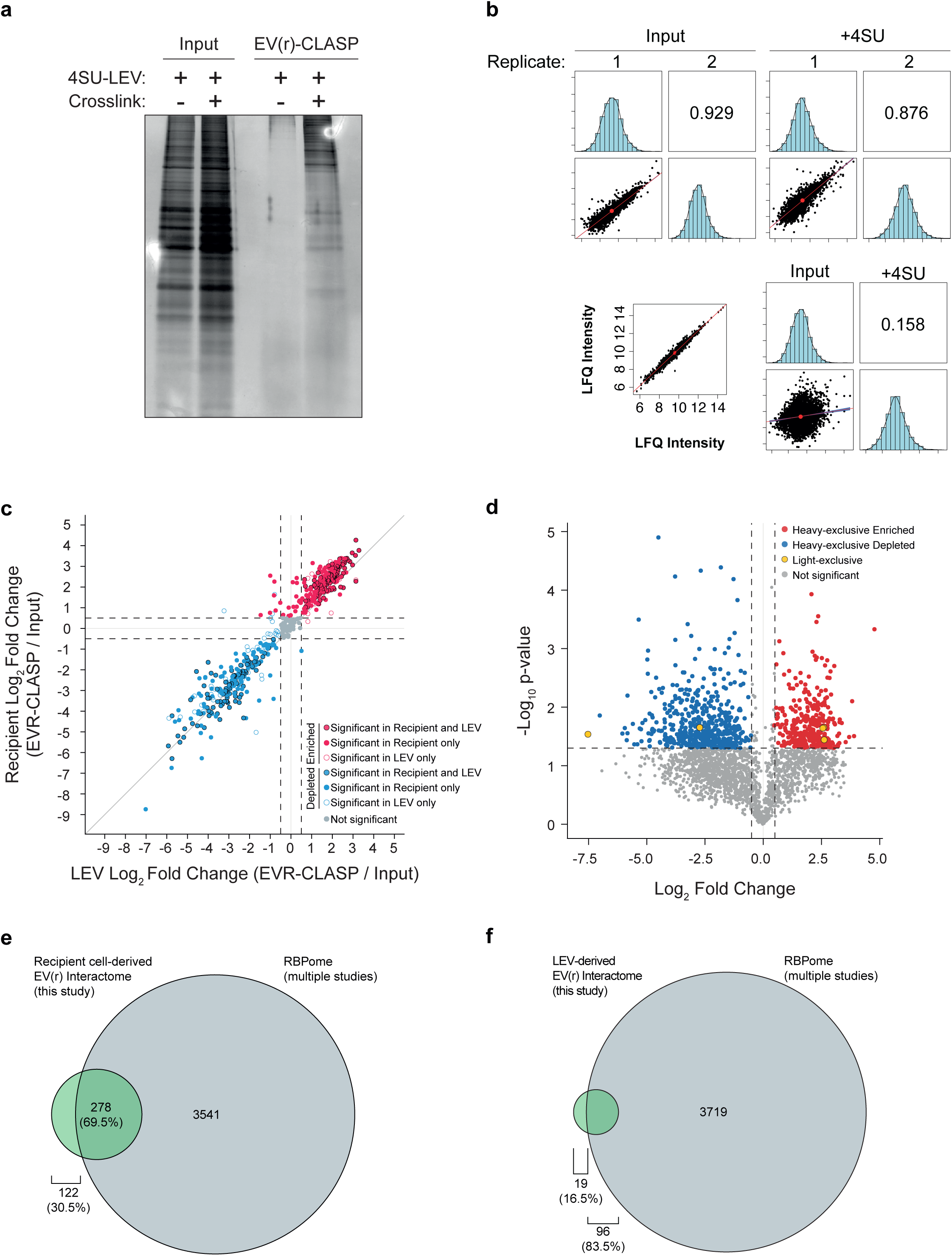
Expected outcome and quantitative proteomic analysis of EV(r)-CLASP samples. **(a)** SDS–PAGE followed by silver staining of proteins enriched by EV(r)-CLASP from 4SU-labeled extracellular vesicles (EVs) isolated from DLD colorectal cancer donor cells following EV uptake by DKs-8 recipient cell lines. Donor cells were cultured in light SILAC medium, whereas recipient cells were maintained in heavy SILAC medium to enable quantitative discrimination between donor-derived EV-associated proteins and recipient-cell-enriched proteins following EV-mediated RNA–protein transfer and regulatory remodeling. **(b)** Pearson correlation coefficient analysis across biological replicate samples of input and EV(r)-CLASP conditions demonstrates strong experimental reproducibility and SILAC labeling consistency (top). Correlation analysis between recipient-cell input and EV(r)-CLASP samples further reveals distinct proteomic enrichment profiles associated with LEV-mediated RNA–protein transfer and recipient-cell remodeling (bottom). **(c)** (c) Comparison of RNA-binding protein enrichment between recipient cells and large extracellular vesicles (LEVs). Each point is one protein, plotted by its log₂ fold change (EVR-CLASP / input) in LEVs (x-axis) and recipient cells (y-axis). Filled circles with black outlines are significant in both datasets, filled circles without outlines are significant in recipient cells only, and open circles are significant in LEVs only. Red points are enriched and blue points are depleted. Grey points are not significant. Dashed lines mark the fold-change cut-off (|log₂ FC| = 0.5), and the solid diagonal line marks y = x. **(d)** Volcano plot of proteins identified by EVR-CLASP in SILAC-labelled recipient cells. Red and blue points are heavy-label-exclusive proteins that are significantly enriched or depleted. Yellow points are light-label-exclusive proteins, which likely come from EVs. Grey points are not significant. The vertical dashed lines mark |log₂ FC| = 0.5, and the horizontal dashed line marks p = 0.05 (−log₁₀ p ≈ 1.3). **(e)** Overlap between the recipient cell–derived EV(r) interactome from this study (green) and well established RNA-binding protein (RBP) reference databases, such as RBP2GO (grey). Of the 400 interactome proteins, 278 (69.5%) are previously annotated RNA-binding proteins and 122 (30.5%) are new candidates. **(f)** Overlap between the LEV-derived EV(r) interactome from this study (green) and the combined RBPome (grey). Of the 115 interactome proteins, 96 (83.5%) are previously annotated RNA-binding proteins and 19 (16.5%) are new candidates.

##### Mass spectrometric analysis ● Timing: ∼7 d

60. Protein samples (10-100 µg) in TEAB buffer are reduced by addition of 20 mM DTT at 95 °C for 10 minutes.

61. Cool to room temperature and alkylate the sample with 40 mM iodoacetamide, incubate in the dark for 30 minutes at room temperature.

62. Acidify the sample by addition of aqueous phosphoric acid (1.2% final).

**CRITICAL STEP:** The acidification of protein allows binding to the S-trap column. Improper preparation at this step will result in loss of sample.

63. Add S-trap Binding Buffer (90% Methanol, 100 mM TEAB, pH 7.5).

64. Load sample mixture onto column and centrifuge 200 xg for 1 minute. Discarding flow-through.

65. Wash column using S-trap Binding Buffer. Centrifuge as before.

66. Repeat wash 3 times.

67. Add trypsin prepared in 50 mM TEAB, pH 8.0 to final concentration of 1 mg/mL or 1:10 to 1:50 trypsin:protein ratio.

68. Incubate for 1 hour at 47 °C.

**!CAUTION** Do not agitate or shake the tubes during trypsin digestion.

69. Sequentially add elution buffers and centrifuge S-trap tubes as before for 1 minute:

(A) Elution 1 – 80 µL 50 mM TEAB (pH 8)

(B) Elution 2 – 80 µL 0.2%-0.5% aqueous formic acid

(C) Elution 3 – 80 µL 50% Acetonitrile, 0.2%-0.5% formic acid

70. Pool eluates and dry under vacuum

71. Desalt peptide samples using C18 spin column before proceeding to LC-MS/MS analysis.

72. Inject up to 1 µg of protein into the LC column and apply an HPLC solvent B gradient optimized for the complexity of the protein sample.

73. Mass spectrometry acquisition settings will depend on the specific instrument platform used; however, these settings generally do not require modification from standard protocols used for protein identification or quantitative proteomics analyses.

##### Proteomic data processing ● Timing: ∼2 h

74. Process DIA LC–MS/MS RAW files using FragPipe with MSFragger for peptide identification and DIA-NN for protein quantification^42,43^.

75. Search spectra against a UniProt FASTA database supplemented with contaminant and decoy sequences. Specify carbamidomethylation of cysteine as a fixed modification and methionine oxidation together with protein N-terminal acetylation as variable modifications.

76. Filter peptide-spectrum matches (PSMs), peptides, and proteins to a 1% false discovery rate (FDR). Export peptide-and protein-level quantification tables for downstream statistical, functional, and bioinformatic analyses.

##### Quantitative analysis ● Timing: ∼2 h

77. Perform quantitative proteomic analysis in RStudio using the MSstats R package.

78. Export peptide intensity values from FragPipe, log2-transform the data, and perform quantile normalization prior to downstream statistical analysis.

79. Evaluate the consistency and reproducibility of biological and/or technical replicates through Pearson correlation analysis and visualization of pairwise sample correlations. (Fig. 2b and Supplementary Fig. 2b).

80. Generate volcano plots using EnhancedVolcano to visualize proteins enriched or depleted relative to the input control. Define significantly enriched or depleted proteins using thresholds of *P* value ≤ 0.05 together with fold-change cutoffs ≥0.5 (Fig. 2c).

##### Functional analysis ● Timing: ∼variable

81. Perform functional analysis of proteomics datasets according to the specific goals and requirements of the experiment to identify enriched biological pathways, molecular functions, and cellular processes associated with differentially abundant proteins. Conduct Gene Ontology (GO), pathway enrichment, and protein–protein interaction analyses to evaluate the functional significance of identified proteins and their involvement in biological networks. Use these analyses to gain biological insight into the molecular mechanisms and cellular pathways associated with the experimental conditions. To further evaluate the recovery of RNA-binding proteins, compare the enriched protein set with established RBP reference databases, such as RBP2GO^44^. A high degree of overlap with annotated RBPs is generally expected, supporting the successful identification of proteins associated with RNA binding and RNA-mediated regulatory pathways (Fig. 2d).

#### Procedure 2: EV(r)-CLASP enables systematic profiling of EV-derived RNA–protein interactions in recipient cells

##### Preparation of SILAC DMEM media ● Timing: ∼30 min

1. When required, supplement the custom-synthesized SILAC DMEM medium (500mL) with standard amino acids in both the light and heavy media.

2. Add the appropriate light (^12^C_6_ ^15^N_2_ L-Lysine and ^12^C_6_ ^15^N_4_ L-Arginine) or heavy (^13^C_6_ ^15^N_2_ L-Lysine and ^13^C_6_ ^15^N_4_ L-Arginine) isotopic forms of lysine and arginine to the corresponding light and heavy SILAC DMEM media.

3. Supplement both media with antibiotics, glutamine, and 10% dialyzed fetal bovine serum.

**PAUSE POINT** SILAC media may be stored at 4 °C for up to 2 months, if needed.

##### Adaptation of Cells from standard DMEM to SILAC medium ● Timing: ∼3-4 weeks

4. Transfer donor (DLD) and recipient (DKs8) cell lines at 80–90% confluency into separate culture dishes containing light and heavy SILAC media, respectively. Seed cells at approximately 15-20% confluency to allow at least two cell doublings in the newly supplemented media.

5. Refresh the light or heavy SILAC medium every 2-3 days if the cells are not yet ready for passage.

6. Perform subculturing of the light and heavy populations in their specific media before they reach total confluency.

7. Begin expanding the labeled cell populations after the first passage by moving them into larger dishes or increasing the number of plates as required.

8. Subculture DLDs and Dks8 cells in the appropriate SILAC medium to allow for a minimum of five cell doublings.

9. Repeat step 8 for at least six passages to ensure sufficient incorporation of SILAC amino acids into both donor and recipient cell lines.

**CRITICAL STEP** Verify that enough time has passed for full amino acid incorporation before moving to the next experimental stage as mentioned in experimental design section. This evaluation is necessary when using a new cell stock to prevent quantification errors caused by incomplete labeling.

**PAUSE POINT** Light- and heavy-labeled cells may be cryopreserved in appropriate freezing medium, such as 10% DMSO in 90% dialyzed FBS, for long-term storage.

##### Generation and isolation of 4SU-labeled EVs from donor cells cultured in light SILAC medium. ● Timing ∼3 d

**CRITICAL STEP** After confirming >95% incorporation of isotopic labels by mass spectrometric analysis of peptides derived from light- and heavy-labeled proteins in donor (DLDs) and recipient (DKs8) cell lines, seed 15 × 10^6^ donor cells in 150 mm tissue culture dishes containing 20 mL complete DMEM supplemented with dialyzed FBS, glutamine, and Pen–Strep.

10. Propagation and isolation of EVs should be done as discussed in Procedure 1, Steps 8 to 28

##### Characterizing EVs using NTA ● Timing ∼2 h

11. Characterize the concentration and size distribution of SILAC labelled donor cell-derived LEV by NTA, as mentioned in box 1, (Supplementary Fig. 4).

#### EV(r)-CLASP

##### Delivery of 4SU-labeled EVs into recipient cells and UV crosslinking ● Timing ∼12 h

12. For each experimental condition, seed 15 × 10^6^ recipient cells in 150 mm tissue culture dishes containing 20 mL complete DMEM (Fig. 1). Typically, 6 plates per experimental condition.

13. Incubate cells overnight at 37 °C in a humidified tissue culture incubator.

14. Remove culture medium and wash cells twice with 5 mL cold PBS.

15. Remove PBS completely and incubate cells with 2.5 mL of diluted 4SU-labeled EV preparation in serum-free medium. Incubate cells at 4 °C for 1 h, with gentle rocking every 10–15 min to ensure even distribution and prevent drying of the cell monolayer

**CRTICAL STEP** The initial EV dose was estimated based on the secretion rate of the parental DLD-1 cells and calculated as the number of EVs secreted per cell per hour multiplied by the number of donor cells and the duration of the assay. This approach was used to approximate the cumulative EV exposure that recipient cells would experience under physiological co-culture conditions.

**?TROUBLESHOOTING**

16. Following incubation, remove the EVs-containing medium (LEVs and SEVs) and wash cells twice with sterile PBS to remove non-internalized extracellular vesicles.

17. Add 17 mL complete DMEM and incubate cells at 37 °C for 10 h.

18. Remove the culture medium and wash cells twice with 5 mL cold PBS.

19. Remove residual PBS and add 2.5 mL cold PBS to each dish.

**CRITICAL STEP** Thorough washing is essential to remove non-internalized EVs and reduce background signals.

20. Irradiate uncovered dishes twice with 1.2 J/cm² of 365 nm UV light using a Spectrolinker XL-1500 or equivalent UV crosslinking device (Fig. 1).

21. Harvest cells using a cell scraper and transfer the cell suspension to 15 mL conical tubes. Centrifuge at 300 × g for 5 min at 4 °C to pellet the cells. Carefully remove and discard the supernatant, then determine the weight of the cell pellet before proceeding with lysis and EV(r)-CLASP purification.

**PAUSE POINT** Snap-freeze cell pellets in liquid nitrogen and store at −80 °C. When ready to continue, resuspend pellets directly in 1× denaturation buffer and mix thoroughly by pipetting.

##### Preparation of Whole-Cell Lysate and Protein Denaturation ● Timing ∼ 5 h

22. Incubate the cells obtained from step 21 in 1 X denaturation buffer (refer to Box 2 for details regarding optimization of cell lysis and SPRI bead:lysate ratios) at 95 °C for 10 minutes. After boiling, transfer two 200 µL aliquots into separate 1.5 mL Eppendorf tubes to serve as “input” samples.

**PAUSE POINT**: Cell lysates may be stored at -20 °C. To resume the protocol, thaw the samples and repeat the 95 °C incubation for 10 minutes.

23. Allow the samples to cool and incubate at RT for 10 minutes.

24. Add SPRI bead suspension (see Reagent Setup) to denatured lysate as mentioned in box 2 resulting in a 2:3 bead suspension-to-sample ratio.

25. Mix the samples by pipetting up and down 10 times, then incubate at 25 °C for 10 min. Place the tubes on a magnetic stand and allow the SPRI beads to collect on the magnet. Bead settling times may vary and can take up to 3 min.

26. Carefully remove and discard the supernatant, then add 1 mL wash buffer.

27. Vortex the samples at maximum speed and allow the SPRI beads to re-pellet on the magnetic stand. Repeat Steps 26 and 27 for three additional times, for a total of four washes.

**? TROUBLESHOOTING**

28. Remove the final wash supernatant, add 600 µL of 1× denaturation buffer, and mix thoroughly by vortexing. Incubate the samples at RT for 5 min.

29. Add 400 µL beads buffer and mix the reaction thoroughly by pipetting 10 times. Incubate the samples at 25 °C for 10 min, then place the tubes on a magnetic stand.

30. Remove and discard the supernatant, then add 1 mL wash buffer.

31. Vortex the samples at maximum speed and allow the SPRI beads to re-pellet on the magnetic stand. Repeat Steps 30 and 31 one additional time for a total of two washes.

32. Place tubes on magnetic stand and allow for all the beads to collect.

33. Remove and discard the supernatant. Resuspend the beads from one tube in 1 mL of wash buffer by vortexing at maximum speed. Transfer the bead suspension to the next tube and repeat the process sequentially to combine the contents of all four sample tubes into a single tube using the same 1 mL wash buffer suspension.

34. Place the tube containing the aggregated beads on a magnetic stand and allow the beads to fully collect.

35. Remove and discard the supernatant and replace it with 1 mL of pre-warmed wash buffer.

36. Vortex the beads at maximum speed to thoroughly resuspend them, then return the tube to the magnetic stand and allow the beads to fully re-pellet.

37. Remove the wash buffer supernatant and add 200 μL of 1× denaturation buffer. Thoroughly resuspend the beads by vortexing.

38. Briefly centrifuge the tubes at low speed for a few seconds to collect beads from the walls and caps of the tubes. Avoid over-centrifugation, as this may cause bead aggregation or clumping.

39. Incubate the samples at room temperature for 5 min.

40. Place the tubes on a magnetic stand to collect the beads and transfer the supernatant to a fresh tube. Save a 20 µL aliquot of the supernatant in a 1.5 mL eppendorf tube for Step 41.

**?TROUBLESHOOTING**

**PAUSE POINT** Eluted EV(r)-CLASP samples at step 40 may be stored at −80 °C for several weeks before Benzonase treatment. Avoid repeated freeze–thaw cycles, as these may reduce sample integrity, protein recovery, and quantitative reproducibility.

41. (Optional) To assess RNA recovery by CLASP, treat the aliquot collected in step 40 with 1 mg/mL proteinase K for 2 h at 55 °C, followed by RNA purification using TRIzol according to the manufacturer’s instructions.

##### Benzonase treatment ● Timing ∼ 2 h

42. Digest nucleic acids and recover crosslinked proteins by Benzonase treatment as described in Procedure 1, steps 46 and 47

##### Methanol and Chloroform Precipitation ● Timing: ∼45 min

43. Perform methanol–chloroform precipitation and protein recovery of EV(r)-CLASP samples as described in Procedure 1, Steps 48-59.

**PAUSE POINT** Protein pellets obtained from EV(r)-CLASP samples may be stored at - 80 °C for several weeks before resuspension and downstream proteomic analysis.

44. Proceed with silver staining to visualize proteins enriched by EV(r)-CLASP samples (Fig. 3a and Supplementary Fig. 3c) or continue directly to LC–MS/MS analysis for identification and quantitative characterization of donor EVs-derived and recipient cell-associated RNA–protein complexes.

##### Mass spectrometric Analysis ● Timing: ∼7 d

45. For mass spectrometric analysis of the protein pellet refer to Procedure 1, steps 60-73.

##### Proteomic Data Processing ● Timing: ∼2 h

46. Launch FragPipe and load the appropriate SILAC workflow for two-channel (Light/Heavy) quantitative proteomic analysis.

47. Import RAW LC–MS/MS files and select UniProt FASTA database for Human Reference Proteome supplemented with contaminant and decoy sequences for false discovery rate (FDR) estimation.

48. In the MSFragger tab, set carbamidomethylation of cysteine (+57.0215 Da) as a fixed modification. Set the following as variable modifications: methionine oxidation (+15.9949 Da), protein N-terminal acetylation (+42.0106 Da) and the heavy SILAC labels ¹³C₆¹⁵N₂-lysine (K8; +8.0142 Da) and ¹³C₆¹⁵N₄-arginine (R10; +10.0083 Da).

49. Select trypsin as the proteolytic enzyme and allow up to two missed cleavages.

50. In the Validation tab, enable MSBooster rescoring with retention time, spectrum and ion mobility prediction (DIA-NN models), and validate peptide spectrum matches with Percolator (minimum probability 0.5). Disable Crystal-C and PTMProphet. Filter results at 1% protein-level FDR (ProteinProphet; picked-protein strategy).

51. In the Spec Lib tab, generate a spectral library from the search results with EasyPQP using psm.tsv as input, automatic selection of reference runs for retention time and ion mobility calibration, b- and y-ion fragments, a UniMod annotation tolerance of 0.02 Da and a fragment annotation tolerance of 15 ppm.

52. In the Quant (DIA) tab, turn on DIA-NN quantification. Set the false discovery rate (FDR) to 1% (0.01), choose a high-precision quantification strategy.

53. Export peptide and protein-level quantification tables for downstream statistical, pathway enrichment, and bioinformatic analyses.

##### Quantitative analysis ● Timing: ∼2 h

54. Import the peptide or protein intensity tables generated by the mass spectrometry data analysis software (e.g., FragPipe) into RStudio and perform quantitative proteomic analysis using appropriate statistical packages, such as the MSstats R package. Normalize the data and estimate protein abundances across all EV(r)-CLASP and control samples.

55. Evaluate the reproducibility of biological and/or technical replicates by calculating Pearson correlation coefficients and visualizing sample-to-sample correlations (Fig. 3b).

56. Analyse differential protein enrichment between EV(r)-CLASP samples and their corresponding inputs, separately for the heavy (recipient-cell) and light (LEV-derived) SILAC channels, classifying proteins with P < 0.05 and |log₂ fold change| > 0.5 as significantly enriched or depleted. For proteins quantified in both channels, plot recipient against LEV log₂ fold changes (Fig. 3c); analyse proteins quantified in only one channel by volcano plot (Fig. 3d). To distinguish recipient-specific from LEV-derived binders, overlay both channels in a single volcano plot (Supplementary Fig. 3d); for example, HNRNPLL and DKC1 are enriched in the recipient channel but not in the LEV channel. Recipient-cell proteins associated with EV-transferred RNAs, together with LEV-derived RNP components, should be enriched.

##### Functional analysis Timing: ∼ variable

57. Perform functional analysis of EV(r)-CLASP datasets to identify biological pathways, molecular functions, and cellular processes associated with extracellular vesicle (EV)-mediated RNA transfer and recipient-cell regulatory remodeling. Perform Gene Ontology (GO) enrichment, pathway enrichment, and protein–protein interaction analyses on significantly enriched recipient-cell-associated and LEV-associated proteins identified from EV(r)-CLASP datasets depending on the researcher’s specific experimental goal. To further evaluate the enrichment of RNA-binding proteins (RBPs), compare the identified protein set with established RBP reference databases, such as RBP2GO; a substantial overlap with annotated RBPs is generally expected and serves as an indicator of successful capture of RNA-associated protein networks (Fig. 3e and 3f).

##### Timing

Procedure 1: EV-CLASP for the systematic identification of extracellular vesicle-associated RNA–protein complexes

Steps 1-7, Propagation of 4SU-labeled EVs: ∼3 d Steps 8-28, Isolation of 4SU-labeled EVs: ∼8 h

Steps 29-30, Characterization of EV purity and concentration by NTA and UV crosslinking: ∼2 h

Steps 31-32, Preparation of EV lysates and protein denaturation: ∼30 min Steps 33-45, Solid-phase purification: ∼3 h

Steps 46-47, Benzonase treatment: ∼2 h

Steps 48-59, Methanol–chloroform precipitation and protein recovery: ∼45 min

Steps 60-73, silver staining or peptide identification and quantification by LC–MS/MS: 3– 5 d depending on number of samples

Steps 74-76, Proteomic data processing: ∼2 h Steps 77-80, Quantitative analysis: ∼2 h

Step 81, Functional analysis: ∼ variable depending on number of goals

Procedure 2: EV(r)-CLASP enables systematic profiling of EV-derived RNA–protein interactions in recipient cells.

Steps 1-3, Preparation of SILAC media: ∼30 min

Steps 4-9, Adaptation of donor and recipient cells to SILAC media: ∼3-4 weeks Step 10, Generation and isolation of 4SU-labeled EVs from donor cells: ∼3 d Step 11, Characterization of EV purity and concentration by NTA: ∼2 h

Steps 12–21, Delivery of EVs into recipient cells and UV crosslinking: ∼12 h Steps 22–41, Preparation of whole-cell lysates and protein denaturation: ∼5 h Step 42, Benzonase treatment: ∼2 h

Step 43, Methanol–chloroform precipitation and protein recovery: ∼45 min

Step 44-45-Silver staining or LC–MS/MS-based peptide identification and quantitative proteomic analysis: ∼3–5 d, depending on sample number and instrument availability.

Steps 46–53, Proteomic data processing: ∼2 h Steps 54–56, Quantitative analysis: ∼2h

Step 57, Functional analysis: ∼ variable depending on number of goals.

##### Anticipated results

Application of EV-CLASP and EV(r)-CLASP is expected to identify robust and reproducible enrichment of EV-RNA-associated protein complexes together with quantitative characterization of recipient-cell RNA regulatory remodeling following EV uptake.

Successful implementation of EV-CLASP should result in efficient recovery of crosslinked EV-associated RNP complexes following UV irradiation and solid-phase purification under denaturing conditions. Following Benzonase treatment and methanol/chloroform precipitation, silver staining or SDS-PAGE analysis should reveal distinct protein bands enriched in 4SU-labeled EV samples relative to negative controls, indicating successful enrichment of RNA-associated proteins and efficient removal of nonspecific contaminants.

NTA of purified EV preparations should demonstrate particle populations within the expected extracellular vesicle size range, typically between 50 and 500 nm depending on the isolation strategy and EV subtype analyzed. EV preparations should display reproducible particle concentrations and narrow size distributions across biological replicates.

LC–MS/MS analysis of EV-CLASP samples is expected to identify hundreds of EV-associated proteins, including known RBPs, translation-associated factors, spliceosomal proteins, stress granule components, RNA helicases, and non-classical RNA-binding proteins. These proteins are anticipated to show significant enrichment in EV-CLASP fractions relative to corresponding input controls and non-4SU-labeled EV controls, demonstrating selective enrichment of EV-associated RNA-protein complexes.

To analyze LC-MS/MS data, use preferred proteomics data analysis software for peptide-spectrum matching and protein identification in consultation with local proteomics experts or core facility personnel. Quantitative proteomic analysis using MSstats should generate highly reproducible protein abundance profiles between biological replicates. Pearson correlation analysis of log2-transformed protein intensities is expected to demonstrate strong reproducibility between replicates, typically with correlation coefficients greater than 0.9. Volcano plot analysis should reveal significant enrichment of EV-associated RNA-binding proteins in EV-CLASP samples relative to input controls, while nonspecific structural and contaminating proteins should remain depleted following stringent denaturing purification.

In EV(r)-CLASP experiments, SILAC-based quantitative proteomics should enable clear discrimination between donor-derived LEV-associated proteins and recipient-cell-associated proteins following LEV uptake. Efficient incorporation of heavy isotopic labels (>95%) should result in robust separation of heavy and light peptide populations during LC-MS/MS analysis, thereby enabling accurate quantitative analysis of EV-mediated RNA-protein transfer and recipient-cell remodeling.

Recipient cells exposed to 4SU-labeled EVs are expected to exhibit significant enrichment of RNA-binding proteins associated with mRNA processing, RNA modification and decay, post-transcriptional regulation, translation control, and intracellular trafficking pathways. Quantitative comparison of EV(r)-CLASP fractions relative to corresponding input controls should identify both donor-derived EV proteins and newly established recipient-cell RNA–protein interactions induced following LEV internalization. Proteins involved in RNA metabolism, ribonucleoprotein assembly, and translational regulation, are expected to display positive log2 fold enrichment values and significant -log10 P values.

Functional enrichment analyses, including GO, pathway enrichment, and protein– protein interaction network analysis, are anticipated to reveal significant enrichment of pathways associated with RNA processing, mRNA stability, translational regulation, intracellular transport, stress response signaling, and EV-mediated intercellular communication. In colorectal cancer models using DLD-1 donor cells and DKs-8 recipient cells, EV(r)-CLASP is expected to identify extensive remodeling of recipient-cell RNA regulatory networks following uptake of oncogenic EV cargo.

Collectively, the EV-CLASP and EV(r)-CLASP methods are anticipated to serve as highly sensitive, scalable, and reproducible platforms for the systematic characterization of EV-associated ribonucleoprotein complexes, EV-mediated RNA transfer, and recipient-cell post-transcriptional regulatory remodeling across diverse biological systems and disease contexts.

#### Reporting Summary

Further information on research design is available in the Nature Research Reporting Summary linked to this article.

#### Data availability

The mass spectrometry proteomics data have been deposited to the ProteomeXchange Consortium^45^ via the PRIDE^46^ partner repository with the dataset identifier PRIDE: *in progress*.

#### Code availability

All code used for proteomic analysis and figure generation is accessible at https://github.com/Ascano-Lab

**Box 1: Particle Tracking for EV analysis Timing ∼2 h**

1. Turn on the ZetaView instrument, laser module, and computer workstation.
2. Launch the ZetaView software and allow the system to initialize.
3. Ensure that the fluorescent filter is removed before proceeding, then click OK.
4. Flush the measurement cell sequentially with deionized water followed by PBS.
5. Prepare a 102 nm standard dilution at 1:250,000 using filtered deionized water.
6. For auto-alignment, load 2–3 mL of the diluted standard into the measurement cell and click OK.
7. Rinse the system using Pump 1 then flush with filtered PBS using a syringe until the detected particle count is below 10.
8. Inspect positions 0.1–0.9 for the presence of bubbles or residual particles. All positions should display particle counts below 10.
9. Load 1 mL of the sample into the measurement cell. The detected particle count should range between 100 and 300. If the count falls outside this range, prepare a new dilution and repeat the procedure starting from step 7.
10. Open the Measurement tab in the ZetaView software.
11. Click Run Video Acquisition and inspect the particle-tracking videos to ensure accurate particle detection and optimal tracking quality.
12. Analyze all recordings using the ZetaView software while maintaining identical post-acquisition analysis settings across all samples.
13. Determine the particle concentration (particles/mL), mean particle diameter, size distribution profile, and particle distribution histograms.
14. Export the raw data and summary statistics for downstream analysis.
15. Following sample acquisition, flush the measurement cell sequentially with particle-free PBS followed by deionized water.
16. Ensure complete removal of residual particles before shutting down the instrument.
17. Transfer both 4SU-labeled EV preparations from the ultracentrifuge tubes into separate 22 cm^2^ Petri dishes.

**Box 2: Optimization of cell lysis and SPRI bead ratio for CLASP**

We have found that the CLASP conditions may require additional optimization depending on the sample type and cell lines used in the study. Generally, if the conditions outlined in the VIR-CLASP protocol^13^ or this protocol does not yield sufficiently low noise within the negative condition by SDS-PAGE and silver stain analysis, the optimization guideline below may provide better sample quality.

1. Determine the approximate amount of total protein present for each mg of wet cell pellet.
2. This is done by collecting a set number of cells by centrifugation (3 million cells)
3. Removal culture medium by sequential washing with PBS and centrifugation.
4. Weigh the cell pellet (ensure a pre-weighed tube is used for the final collection in step 3).
5. Lyse the cells in 1X denaturation buffer and determine total protein by BCA.
6. Calculate the amount of 1X denaturation buffer to lyse each mg of cell pellet to yield a protein concentration between 0.4-0.6 mg/mL. For example: For DKs-8 cells 45 mg (3 million cells) of wet cell pellet contains approximately 1.3 mg total protein. This equates to 0.03 mg protein/mg wet pellet. Therefore, to obtain 0.4-0.6 mg/mL final protein concentration, each mg of wet cell pellet must be lysed using 50-75 µL 1X denaturation buffer.
7. Multiply the wet cell pellet weight by the 1X denaturation buffer calculated above:

For example: 100 mg wet pellet x 55 = 5500 µL 1X denaturation buffer. Removing 200 µL as input samples (x2) leaves a remaining 5100 µL lysate. To this 0.66x ratio of SPRI beads is added (for volume calculation divide 5100 by 1.5) – thus 3600 µL SPRI bead suspension is added to the lysate for a final 9100 µL lysate-bead mixture.

## Acknowledgements

We thank K. L. Rose and the Vanderbilt Mass Spectrometry Research Center for processing the mass spectrometry samples. We also thank the members of the Ascano laboratory for their support, collegiality, and critical review of the manuscript. This work was supported by the National Science Foundation under Award No. MCB-2330665 (MA,AW), R35 Award No. 5R35GM119569 (MA), and by the National Institute of Diabetes and Digestive and Kidney Diseases (NIDDK) under Award Nos. R01DK123181 (TN, MA) and R01DK134646 (TN, MA). This work was also made possible, in part, through access to shared resources, state-of-the-art instrumentation, and the collaborative research environment provided by the Vanderbilt Center for Extracellular Vesicle Research.

## Author contributions

T.N. and M.A. initiated the concept and design. R.D., S.C., Y.S., A.W., and B.B. further developed design and conceptualization. R.D., S.C., M.K., M.G. II., B.B., and Y.S. optimized the EV-CLASP and EV(r)-CLASP protocols. R.D. and S.C. performed proteomic analysis and data visualization. R.D., S.C., and M.A. wrote the paper with editing by all coauthors.

## Additional information Related links

### Key references using this protocol

Kim, B., et al. (2020). "Discovery of Widespread Host Protein Interactions with the Pre-replicated Genome of CHIKV Using VIR-CLASP." Mol Cell 78(4): https://doi.org/10.1016/j.molcel.2020.04.013

Barrows, N. J., et al. (2019). "Dual roles for the ER membrane protein complex in flavivirus infection: viral entry and protein biogenesis." Sci Rep **9**(1): 9711. https://doi.org/10.1038/s41598-019-45910-9

Kim, B., et al. (2021). "Viral crosslinking and solid-phase purification enables discovery of ribonucleoprotein complexes on incoming RNA virus genomes." Nat Protoc **16**(1): 516–531. https://doi.org/10.1038/s41596-020-00429-6

