## Supplementary Files for "Extracellular vesicle cross-linking coupled with solid-phase purification facilitates discovery of ribonucleoprotein complexes assembled on EV RNAs"

### Supplementary Figure 1.

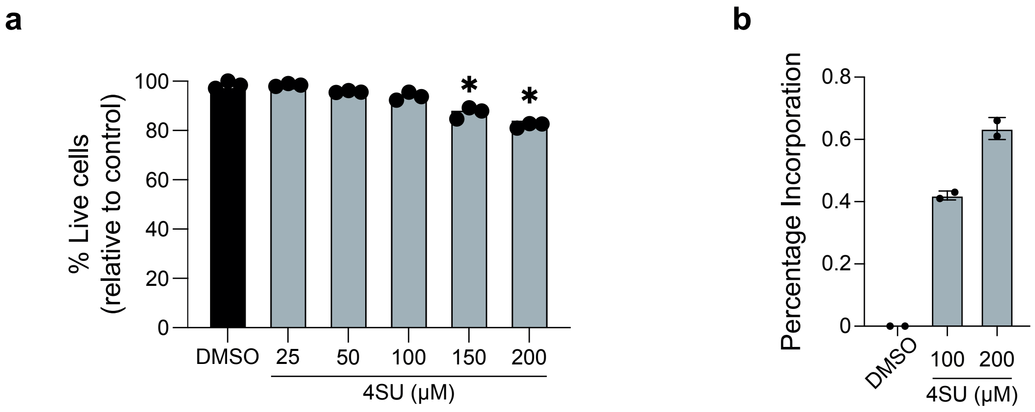


**Supplementary Figure 1. 4-Thiouridine incorporates into DLD-1 RNA without loss of viability.** DLD-1 cells were cultured in conditioning medium supplemented with 4SU. (a) The cell viability monitored after supplementation with 25-200 µM 4SU and (b) the percentage of 4SU incorporation into total RNA was determined by RP-HPLC as described previously[40].

### Supplementary Figure 2.

**a**

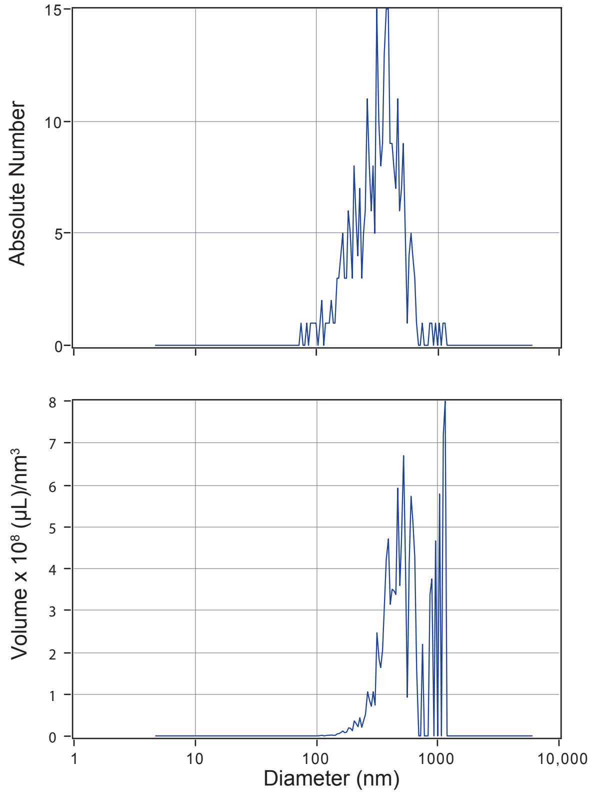


##### Result

| | Number | Concentration | Volume |
|---|---|---|---|
| Median (x50) | 329.1 | 329.1 | 519.4 |
| Std. Deviation | 161.0 | 160.8 | 275.8 |

| | |
|---|---|
| Concentration | $4.9 \times 10^7$ Particles/mL |
| Dilution Factor | 250 |
| Original Concentration | $1.2 \times 10^{10}$ Particles/mL |

##### Quality

| | |
|---|---|
| Ave Particles/Frame | 137 |
| Traced Particle Count | 309 |

##### Peak Analysis

| Diameter (nm) | Absolute Number | FWHM (nm) | Percentage |
|---|---|---|---|
| 379.6 | $1.4 \times 10^1$ | 220.0 | 67.6 |
| 210.2 | $6.0 \times 10^0$ | 68.4 | 22.6 |
| 270.2 | $8.3 \times 10^0$ | 53.8 | 9.7 |

##### X Values

| | Number | Concentration | Volume |
|---|---|---|---|
| X10 | 164.9 | 164.9 | 327.5 |
| X50 | 329.1 | 329.1 | 519.4 |
| X90 | 519.9 | 519.9 | 1086.9 |
| Span | 1.1 | 1.1 | 1.5 |
| Mean | 350.5 | 350.5 | 621.1 |
| Std. Deviation | 161.0 | 160.8 | 275.8 |

**b**

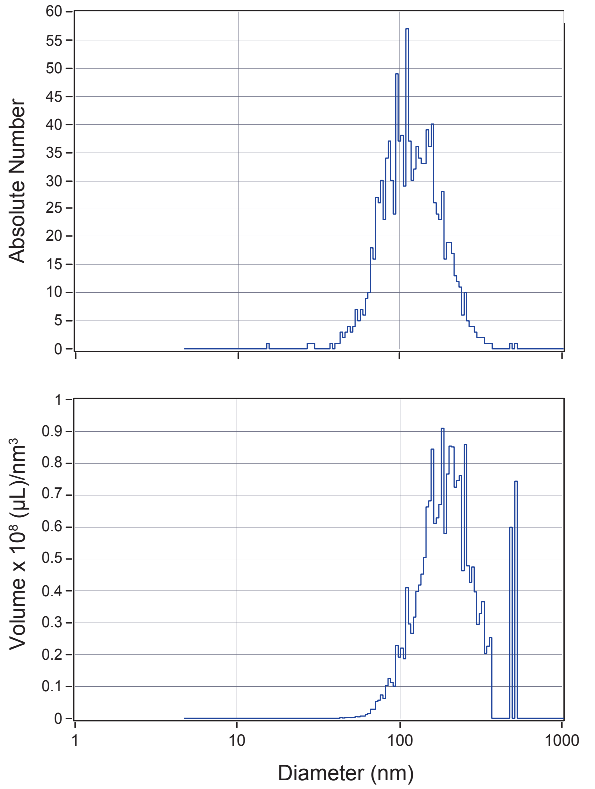


##### Result

| | Number | Concentration | Volume |
|---|---|---|---|
| Median (x50) | 114.1 | 114.1 | 192.9 |
| Std. Deviation | 54.8 | 54.8 | 99.5 |

| | |
|---|---|
| Concentration | $9.1 \times 10^7$ Particles/mL |
| Dilution Factor | 40,000 |
| Original Concentration | $3.6 \times 10^{12}$ Particles/mL |

##### Quality

| | |
|---|---|
| Ave Particles/Frame | 253 |
| Traced Particle Count | 1146 |

##### Peak Analysis

| Diameter (nm) | Absolute Number | FWHM (nm) | Percentage |
|---|---|---|---|
| 110.0 | $4.2 \times 10^1$ | 53.3 | 56.4 |
| 150.9 | $3.7 \times 10^1$ | 68.7 | 43.6 |

##### X Values

| | Number | Concentration | Volume |
|---|---|---|---|
| X10 | 70.6 | 70.6 | 112.0 |
| X50 | 114.1 | 114.1 | 192.9 |
| X90 | 197.7 | 197.7 | 324.2 |
| Span | 1.1 | 1.1 | 1.5 |
| Mean | 128.5 | 128.5 | 215.9 |
| Std. Deviation | 54.8 | 54.8 | 99.5 |

### Supplementary Figure 2 (continued).

**c**

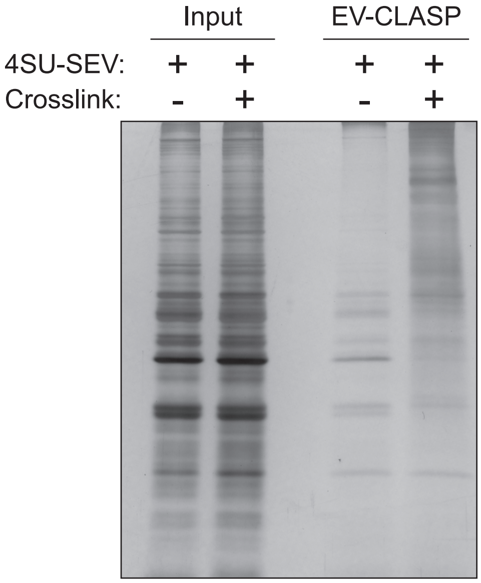


**Supplementary Figure 2. EV characterization by NTA and SEV EV-CLASP.** Particle concentration and size distribution of isolated a) LEVs and b) SEVs were determined by NTA. c) SDS-PAGE and silver stain demonstrating enrichment of RBPs from EV-CLASP.

### Supplementary Figure 3.

**a**

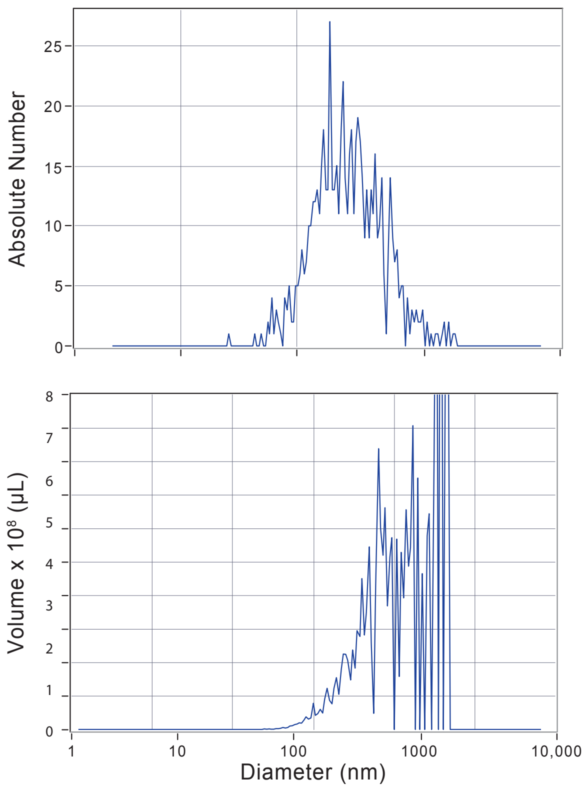


##### Result

| | Number | Concentration | Volume |
|---|---|---|---|
| Median (x50) | 226.9 | 226.9 | 748.8 |
| Std. Deviation | 190.8 | 190.7 | 387.6 |

| | |
|---|---|
| Concentration | 7.6 x $10^{7}$ Particles/mL |
| Dilution Factor | 250 |
| Original Concentration | 1.9 x $10^{10}$ Particles/mL |

##### Quality

| | |
|---|---|
| Ave Particles/Frame | 211 |
| Traced Particle Count | 656 |

##### Peak Analysis

| Diameter (nm) | Absolute Number | FWHM (nm) | Percentage |
|---|---|---|---|
| 176.0 | 1.8 x $10^{1}$ | 61.5 | 34.5 |
| 281.3 | 1.7 x $10^{1}$ | 58.3 | 19.9 |
| 219.1 | 1.8 x $10^{1}$ | 22.9 | 14.1 |
| 372.9 | 1.3 x $10^{1}$ | 81.1 | 13.3 |
| 499.1 | 1.0 x $10^{1}$ | 89.1 | 10.4 |

##### X Values

| | Number | Concentration | Volume |
|---|---|---|---|
| X10 | 116.7 | 116.7 | 321.6 |
| X50 | 226.9 | 226.9 | 748.8 |
| X90 | 494.2 | 494.2 | 1346.9 |
| Span | 1.7 | 1.7 | 1.4 |
| Mean | 284.0 | 284.0 | 814.8 |
| Std. Deviation | 190.8 | 190.7 | 387.6 |

**b**

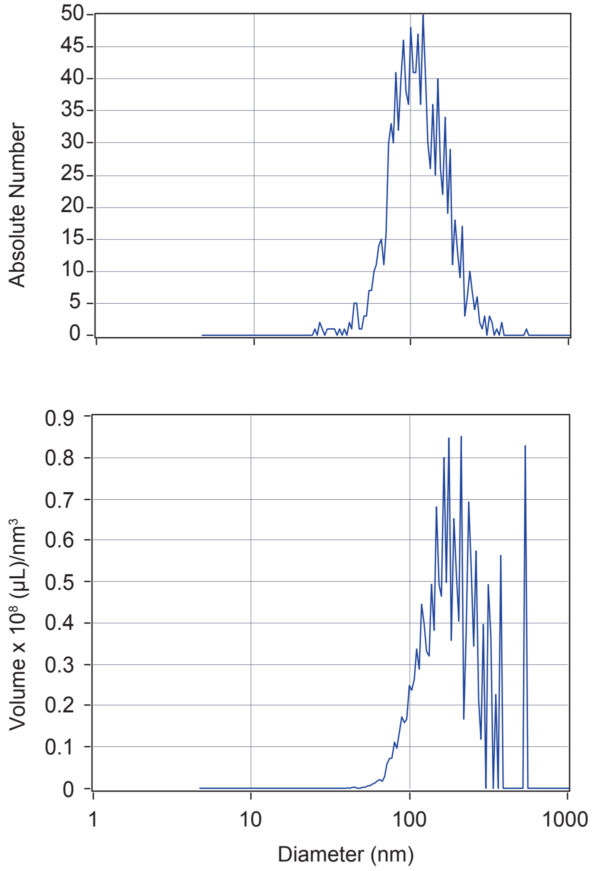


##### Result

| | Number | Concentration | Volume |
|---|---|---|---|
| Median (x50) | 107.2 | 107.2 | 177.2 |
| Std. Deviation | 51.0 | 50.9 | 103.7 |

| | |
|---|---|
| Concentration | 1.0 x $10^{8}$ Particles/mL |
| Dilution Factor | 4000 |
| Original Concentration | 4.1 x $10^{11}$ Particles/mL |

##### Quality

| | |
|---|---|
| Ave Particles/Frame | 282 |
| Traced Particle Count | 1157 |

##### Peak Analysis

| Diameter (nm) | Absolute Number | FWHM (nm) | Percentage |
|---|---|---|---|
| 113.1 | 4.3 x $10^{1}$ | 62.1 | 68.1 |
| 143.3 | 3.2 x $10^{1}$ | 50.1 | 30.0 |
| 44.1 | 3.8 x $10^{0}$ | 3.9 | 1.9 |

##### X Values

| | Number | Concentration | Volume |
|---|---|---|---|
| X10 | 68.6 | 68.6 | 104.4 |
| X50 | 107.2 | 107.2 | 177.2 |
| X90 | 179.2 | 179.2 | 323.0 |
| Span | 1.0 | 1.0 | 1.2 |
| Mean | 119.7 | 119.7 | 207.6 |
| Std. Deviation | 51.0 | 50.9 | 103.7 |

### Supplementary Figure 3 (cont.).

c

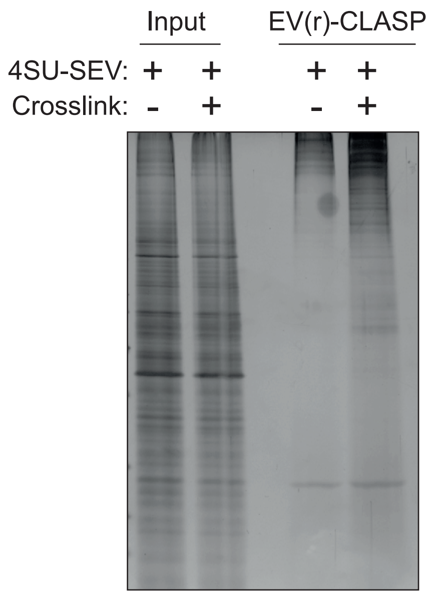


d

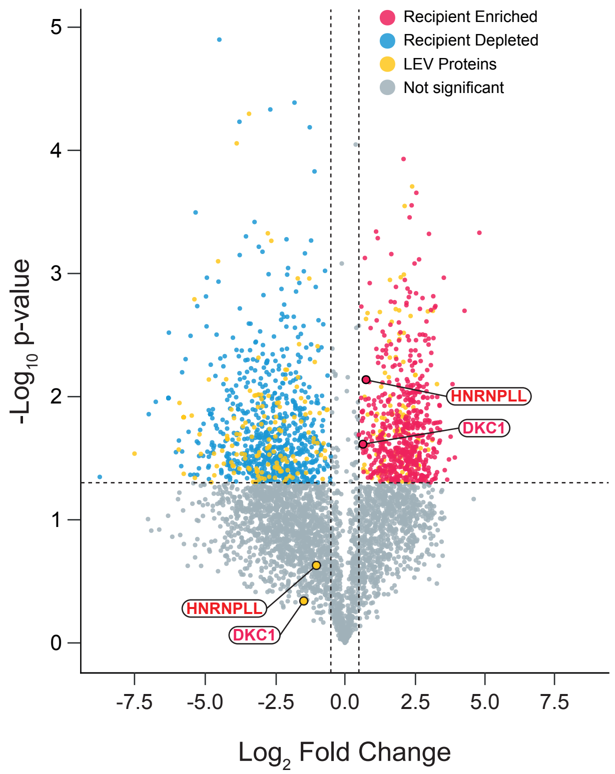


**Supplementary Figure 3. Characterization of SILAC-labelled EVs by NTA for EV(r)-CLASP and SEV EV(r)-CLASP.** Particle concentration and size distribution of EVs isolated from SILAC-labelled donor cells were measured by NTA for a) LEV and b) SEV. c) SEV EV(r)-CLASP silver stain demonstrating enrichment of proteins in 4SU-SEV samples. d) Volcano Plot of SILAC-labelled LEVs processed through the EV(r)-CLASP protocol and DIA proteomics analysis provided protein identifications from heavy and light channel peptides.
